# Mitochondrial inheritance as an important parameter for the rational design of synthetic yeast polyploids

**DOI:** 10.64898/2026.09.22.753424

**Authors:** Sara Orellana-Muñoz, Chloé Haberkorn, Bálint Csoboz, Rike Stelkens, David Peris

## Abstract

Synthetic allopolyploidization is a powerful strategy for generating strains with novel industrially relevant traits. During the design of select, hybridize, evolved and learn cycle, the mitochondrial inheritance is considered neutral and most studies overcome their phenotypic impact. Here, we generated 168 synthetic allotetraploid strains representing all pairwise parental combinations of eight wild *Saccharomyces* species, and systematically controlled mitochondrial inheritance in these crosses. These allopolyploids were phenotyped across 31 environmental conditions relevant to biotechnology. In ∼50% of all condition-strain combinations conditions, allotetraploids exhibited mid-parent heterosis. In approximately 25% of conditions, we found crosses with hybrid vigor, particularly in combinations involving *Saccharomyces arboricola* x *Saccharomyces mikatae*. However, allotetraploids were highly unstable and rapidly sporulated. Our analyses revealed that environmental conditions accounted for the largest effects, follow by its interaction with mitotype. However, mitotype alone showed small effects but significantly affected 42% of the tested conditions, underscoring the importance of mitochondrial inheritance as a key determinant of allopolyploid performance. These findings demonstrate that mitochondrial inheritance is a key, environment-dependent determinant of allopolyploid fitness and should be considered alongside nuclear genome composition in the rational design of industrial allopolyploid strains.

## Introduction

Allopolyploidization naturally occurs across a wide range of eukaryotes, combining complete chromosome sets from distinct species into a single genome. This generates immediate genetic novelty by increasing heterozygosity and gene redundancy, which can buffer deleterious mutations and facilitate adaptive flexibility in fluctuating environments (Comai, 2005). Allopolyploid organisms can display heterosis or hybrid vigor, characterized by enhanced growth rates, stress tolerance, and reproductive success compared to parents. These novel phenotypes may facilitate the occupation of ecological niches that are inaccessible or suboptimal for either parental species (Soltis and Soltis, 2009). Understanding these processes has been instrumental for crop improvement, both through natural selection and induced polyploidization. Many important agricultural species are allopolyploids. For example, the domestication success of wheat (*Triticum aestivum*), cotton (*Gossypium hirsutum*), and canola (*Brassica napus*) has been partly attributed to allopolyploidization, which has generated genetic and phenotypic variation associated with higher yield, stress tolerance, and disease resistance (Chen, 2007; Van de Peer *et al*., 2017). Polyploidization is also observed in animals. Some frog species (e.g., *Xenopus laevis*) display polyploidy, which has been linked to increased environmental tolerance, hybrid vigor, and species diversification, allowing them to colonize diverse ecological niches (Otto, 2007). These examples highlight the potential of allopolyploid-based strategies as a complement to genetic engineering and classical breeding techniques.

In microbial eukaryotes, over the past few decades, multiple strategies have emerged to optimize yeast strains for targeted applications. One accessible avenue involves exploring the untapped potential of wild yeast biodiversity (Peter *et al*., 2018; Libkind *et al*., 2020; Shen *et al*., 2020; Peris *et al*., 2023). Although only a small portion of naturally occurring yeasts has been identified and studied, researchers aim to further enhance these organisms through artificial means. While genetic engineering has been the predominant approach since the 1980s, there has been a renewed interest in classical methods that utilize wild-type strains. Recent studies highlight the advantages of revisiting these strategies, which capitalize on the adaptive potential of naturally isolated yeasts (Stelkens and Bendixsen, 2022; Villarreal *et al*., 2025). One such approach is allopolyploidization, in which complete genomes from different species are combined through the fusion of genetically distinct cells, generating allopolyploid lineages with novel and advantageous phenotypes (Krogerus *et al*., 2015; Krogerus, Magalhães, *et al*., 2017; Peris, Moriarty, *et al*., 2017; Peris *et al*., 2020). Genomic analyses have shown that interspecific genome merging, i.e. hybridization, is not rare among wild *Saccharomyces* isolates (Leducq *et al*., 2017; Peris, Arias, *et al*., 2017) and even more prominent in industrial environments (Gallone *et al*., 2019; Langdon *et al*., 2019), likely as a result of adaptation to the harsh conditions. Currently, nine naturally occurring species are recognized within the *Saccharomyces* genus: *S. arboricola*, *S. cerevisiae*, *S. kudriavzevii*, *S. mikatae*, *S. paradoxus*, *S. jurei*, *S. eubayanus*, *S. uvarum*, and *S. chiloensis* (Desmazières, 1827; Beijerinck, 1898; Bachinskaya, 1914; Naumov *et al*., 2000; Wang and Bai, 2008; Libkind *et al*., 2011; Naseeb *et al*., 2017; Peña *et al*., 2024). Allopolyploids between these species often undergo genomic rearrangements and show single-nucleotide mutations (SNPs) and gene copy number variations (CNVs), which can affect protein complex formation and cellular function (Leducq *et al*., 2012; Peris, Belloch, *et al*., 2012; Piatkowska *et al*., 2013; Hewitt *et al*., 2014; Dandage *et al*., 2021). These genomic changes can generate novel phenotypes through the formation of unique allelic combinations, including heterosis, which can improve the performance of allopolyploid strains in biotechnological applications under certain genetic and environmental conditions (Naseeb *et al*., 2021).

Beyond nuclear genomic contributions, mitochondrial genotype, or mitotype, has been shown to significantly affect allopolyploid performance. Although the nuclear genome encodes the majority of cellular functions, mitochondrial DNA (mtDNA) carries essential components for energy metabolism, particularly those involved in the respiratory chain and mitochondrial gene expression (Adams, 2003; Malina *et al*., 2018). Mitochondria are crucial for ATP production in most eukaryotes. However, in *Saccharomyces* yeasts, mitochondrial function is not strictly essential, as cells can survive through fermentation even in the absence of fully functional mitochondria, as observed in *rho^-^* (partially defective mtDNA) and *rho^0^* (complete loss of mtDNA) strains (Sherman, 1963). Despite this metabolic flexibility, mitochondrial activity depends on tight coordination between mitochondrial and nuclear genomes, involving approximately 750 nuclear-encoded proteins that interact with mitochondrial gene products (Sickmann *et al*., 2003). Accordingly, proper cellular function requires the close coevolution of both genomes (Burton and Barreto, 2012; Piccinini *et al*., 2021). In *Saccharomyces*, mitochondrial inheritance is biparental, resulting in an initial heteroplasmic state that typically resolves within 20–25 generations, leading to fixation of a single mitotype and the establishment of homoplasmy (Strathern *et al*., 1981). Importantly, mitotype inheritance has been associated with functional and evolutionary consequences. For example, all known *S. pastorianus* strains (i.e. allopolyploids between *S. cerevisiae* and *S. eubayanus*) have retained the mtDNA of their cryotolerant parental species *S. eubayanus* (Peris *et al*., 2014; Peris, Arias, *et al*., 2017; Langdon *et al*., 2019). This pattern extends to other industrial allopolyploid yeast strains, where non-*S. cerevisiae* mitotypes are often favored (Peris, Belloch, *et al*., 2012; Peris *et al*., 2018; Langdon *et al*., 2019; Bendixsen *et al*., 2021). Such trends are likely driven by the contribution of mtDNA to key adaptive traits, including low-temperature tolerance, which is critical in lager brewing and certain wine fermentations (Baker *et al*., 2019; Li *et al*., 2019). In addition to temperature adaptation, mitochondrial inheritance can modulate fitness across diverse environmental contexts, including variation in carbon sources (Molinet *et al*., 2024), as well as influence nuclear genome retention and stability in hybrids (Peris, Lopes, *et al*., 2012; Peris *et al*., 2020). Together, these findings highlight mitochondrial genetics as a key and often underappreciated determinant of allopolyploid phenotype, challenging a purely nuclear view of genotype–phenotype relationships and emphasizing the central role of cytonuclear interactions in shaping phenotypic outcomes.

Despite growing evidence highlighting the role of mitochondrial inheritance in industrial *Saccharomyces* allopolyploids, most recent studies (Gyurchev *et al*., 2022; Haberkorn *et al*., 2026; Martínez and Lang, 2026; Rinta-Harri *et al*., 2026) have not systematically controlled for or explicitly considered the impact of mitotype when interpreting fitness outcomes. This gap limits our understanding of how mitochondrial variation shapes allopolyploid performance. Here, we hypothesize that mtDNA plays an important, environment-dependent role in determining the fitness of synthetic *Saccharomyces* allotetraploids, significantly influencing performance in industrially relevant conditions. To test this, we generated a comprehensive collection of 168 allotetraploid strains (56 nuclear-mitotype combinations x 3 replicates) using eight of the nine known diploid *Saccharomyces* species. We then systematically evaluated their fitness across 31 conditions, a subset of industrially-related processes, including multiple stresses, temperature, nutrient limitations, and pH variation. Our results demonstrate that both nuclear and mitochondrial genomes jointly shape allopolyploid fitness, underscoring mitochondrial inheritance as an important, and often overlooked, determinant of allopolyploid performance.

## Experimental procedures

### Yeast strains and maintenance

Our species representative strains of *S. cerevisiae*, *S. paradoxus*, *S. mikatae*, *S. kudriavzevii*, *S. arboricola*, *S. jurei*, *S. uvarum*, and *S. eubayanus* were selected based on previous phenotypic traits of industrial interest (Supplementary Table 1) (Peris, Moriarty, *et al*., 2017; Naseeb *et al*., 2018; Peris *et al*., 2023). *Saccharomyces chiloensis* was not included in this study because it had not yet been described as a distinct species when the work was conducted (Peña *et al*., 2024).

Laboratory strains of *S. cerevisiae*, including a haploid and a diploid version of S288C (Supplementary Table 1), were used as controls. Yeast strains were stored in YPD (1% yeast extract, 2% peptone, 2% glucose) and 15% glycerol at −80 °C.

### Yeast transformation with iHyPr plasmids

Diploid *Saccharomyces* strains of each eight species were transformed with one of three versions of iterative Hybrid Production (iHyPr) plasmids before further manipulation, to enable inducible mating-type switching and subsequent hybridization (Figure 1, Supplementary Table 2). Transformations were performed using the lithium acetate/PEG-4000/carrier DNA protocol, with temperature adjustments tailored to the thermal tolerance of each species (Alexander *et al*., 2016).

**Figure 1.**
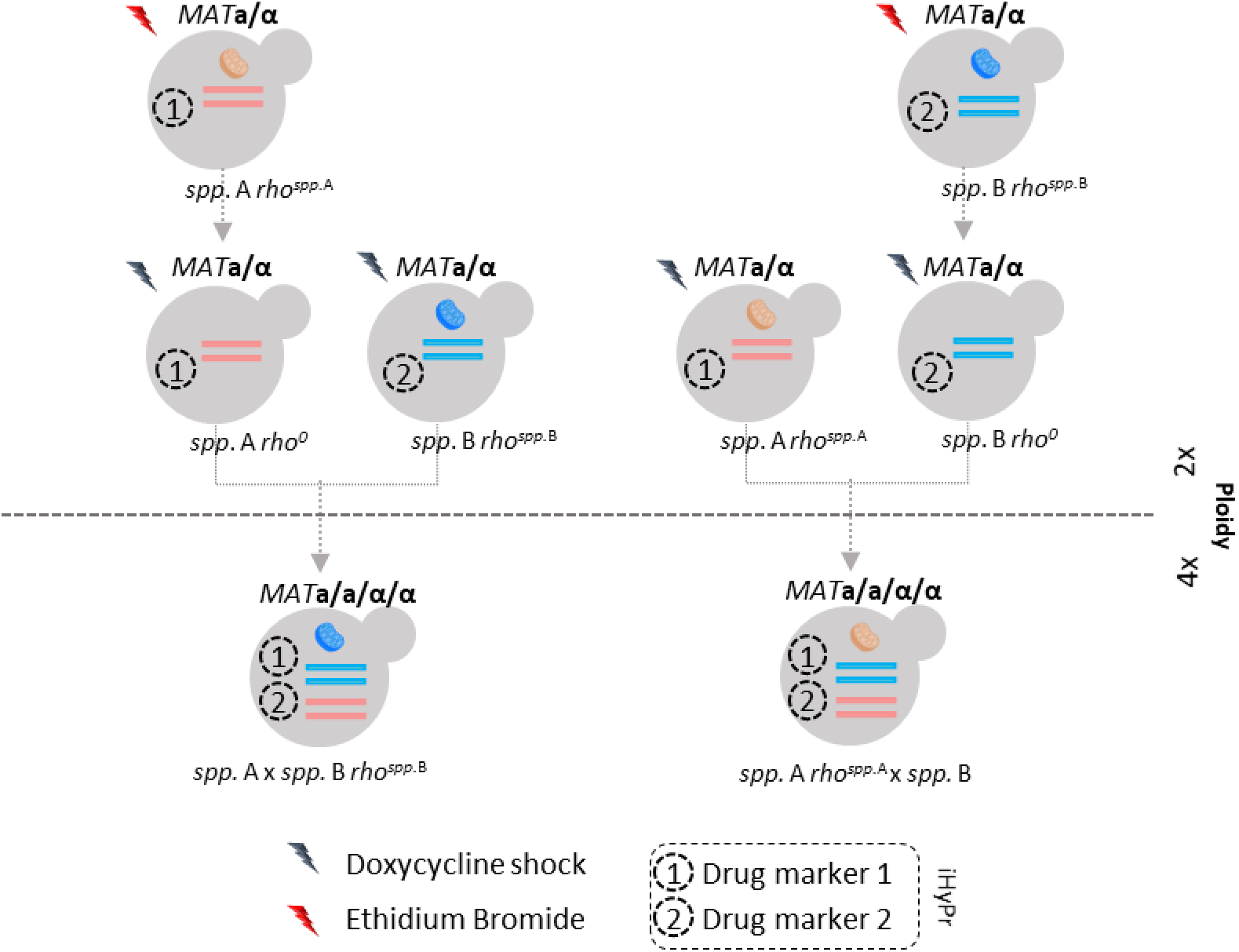
Scheme of the generation of synthetic allotetraploids. Synthetic allotetraploids were generated using the iHyPr and mitoFIX molecular methods (Alexander *et al*., 2016; Baker *et al*., 2019; Peris *et al*., 2020). Dashed arrows indicate hybridization steps. Dashed-line circles represent the iHyPr plasmids, which can carry either drug marker 1 or drug marker 2. The information for each species (*spp*.) or allotetraploid regarding mitochondrial inheritance is noted as either "*rho*" (superscript *spp*. A or *spp*. B) or "*rho^0^*" (lacking mitochondrial DNA).

### Rho^0^ strain generation (rhozerization) with mitoFIX method

The generation of *rho^0^*, strains lacking mitochondrial DNA, was conducted following the mitoFIX method (Baker *et al*., 2019). Strains containing the iHyPr plasmids were grown to saturation in minimal medium (MM) containing 2% glucose and the appropriate antibiotic (Nourseothricin, 100 µg/mL; G418, 50 µg/mL; or Hygromycin B, 200 µg/mL). Cultures were diluted 1:10 into MM supplemented with 25 µg/mL ethidium bromide (EtBr) and incubated to saturation. Cultures were streaked onto YPDA with drug to isolate single colonies. Complete loss of mitochondrial DNA was confirmed by the inability to grow on YPG (glycerol) medium and by the absence of mitochondrial DNA staining following DAPI visualization (Eckert-Boulet *et al*., 2011). Growth on YPG was used to discriminate *rho*⁺ (overnight growth), *rho*⁻ (slow growth in 2 days), and *rho^0^* (no growth) phenotypes (Supplementary Figure 1A). For DAPI staining, cells were grown to exponential phase in MM, diluted to OD₆₀₀ ∼0.2, stained with 1 µg/mL DAPI, incubated at 30 °C for 30 min in darkness, and washed prior to microscopic examination (Supplementary Figure 1B). The strains *rho^+^* and *rho^0^* with iHyPr plasmids were used as parental lines for the generation of two-species allotetraploids with different mtDNA versions.

### Generation of allotetraploids with iHyPr method

A total of 168 allotetraploids (28 diploid nuclear combinations x 2 mitochondrial types x 3 replicates) were generated following the previously described iHyPr method (Alexander et al., 2016; Peris et al., 2020; Ramírez-Aroca, 2019). *rho^+^* and *rho^0^* parental diploid strains with iHyPr plasmids were cultured in YPD containing doxycycline at room temperature overnight (Supplementary Figure 2A). Doxycycline-induced expression of the *HO* endonuclease promoted mating between diploid strains by affecting the *MAT**a***/*MAT**α*** loci (Baker *et al*., 2019). Equal volumes, 10 µl of each parent strain, were mixed and plated onto YPD for 2-3 days at the lowest optimum temperature of both strains, and then a streak on YPD supplemented with appropriate antibiotics to select for successful hybridization (allopolyploidization) events.

### PCR-RFLP confirmation of allopolyploidization

Allopolyploid identity was validated by PCR–restriction fragment length polymorphism (PCR-RFLP) analysis. Genomic DNA was extracted using the phenol:chloroform method from cultures grown to saturation in YPD. Cell lysis was performed in microcentrifuge tubes containing acid-washed glass beads, followed by addition of lysis buffer (10 mM Tris-HCl pH 8.0, 1 mM EDTA, 100 mM NaCl, 1% SDS, 2% Triton X-100) and phenol:chloroform (1:1). Samples were vortexed for 3–4 min and centrifuged at 21,130 × g for 5 min. The aqueous phase was precipitated with 100% ethanol at −80 °C for 10–15 min, washed with 70% ethanol, and resuspended in 100 µL Elution Buffer (EB) at 50–60 °C. RNA was removed by treatment with RNase A (0.5 µL of 10 mg/mL) for 30 min at 37 °C. Genes *BRE5* and *GAL4* were amplified by PCR using newly designed primers following conditions described in González et al., 2006 (Supplementary Table 3). PCR products, confirmed by 1.5% agarose gel, were digested using restriction enzymes capable of distinguishing between *Saccharomyces* species (Peris, Belloch, *et al*., 2012; Alexander *et al*., 2016). Species-specific discriminatory bands were detected by 3% agarose gel electrophoresis (Supplementary Figure 2B).

### Mitochondrial DNA restriction profiling

Mitochondrial DNA profiles were analyzed by restriction digestion. Cells were grown to saturation in YPD, harvested by centrifugation, and resuspended in EB buffer (10 mM Tris-HCl, pH 8.0) containing Zymolyase 20T (8 µL of 5 mg/mL). Spheroplasts were incubated at 37 °C for 30 min, followed by lysis with DNA buffer and phenol:chloroform. After vortexing and centrifugation, the aqueous layer was precipitated with ethanol, washed, and resuspended in Milli-Q water. DNA was digested with *Hinf*I (Boehringer, Mannheim, Germany) and incubated at 37 °C overnight. Restriction patterns were resolved on 1% agarose gels in TAE buffer (40 mM Tris-acetate, 1 mM EDTA, pH 8.0, SafeRed 1×) and visualized by UV transillumination (Supplementary Figure 2C).

### Ploidy estimation by flow cytometry

Cultures were grown to saturation and diluted 1:200 in fresh medium. Once Optical Density at 600 nm (OD₆₀₀) reached 0.4–0.6, aliquots were fixed in 70% ethanol overnight at 4 °C, then treated with RNase A and Proteinase K. Nuclei were stained with SYTOX Green dye (Molecular Probes) (Haase and Reed, 2002). Samples were sonicated and analyzed on an Attune NxT flow cytometer (Invitrogen) using a 488 nm BL1 laser. Green fluorescence was collected at 523 nm. Flow cytometry data were analyzed in FlowJo v10.4.2 (FlowJo, 1996) using the Watson (Pragmatic) model to determine G1 and G2 DNA content. Ploidy levels were estimated by comparison with reference haploid and diploid S288C strains (Supplementary Table 1, Supplementary Figure 2D). DNA content and cell size (FCS-A) correlation analyses was performed using Spearman’s rank-sum test in `R` (R Development Core Team, 2010). Visualizations were generated using `ggplot2` (Wickham, 2009).

### Sporulation confirmation

To evaluate sporulation, cells were growth on YPD medium in agitation over night at 22°C, and washed with dH_2_O. Differential Interference Contrast (DIC) images were acquired on a Carl Zeiss Axioplan 2 imaging microscope using a 63X/1.40NA immersion objective. The microscope was connected to a AxioCam HRc camera controlled by `Carl Zeiss MTB2004 Application Suite` software.

### Phenotypic characterization

The newly generated allotetraploid collection (Supplementary Table 1) was phenotypically assessed in two independent biological replicates using 96-well plates (plates 1–3). Strains were exposed to a wide range of stress conditions and evaluated for multiple phenotypic traits as previously done in Ramírez-Aroca, 2019, including temperature stress (4 °C, 10 °C, 30 °C, and 37 °C), ethanol tolerance (10%, 12%, and 15%), pH (pH 3.5, 7.5, and 10), as well as two different types of metabolic stress, including oxidative and osmotic stress. The latter included exposure to 4 g/L acetate, <50 µM Fe (depleting Fe with BPS), 7 mM and 3 mM FeCl₃, 4 mM potassium metabisulfite, 6 mM and 1 mM sodium sulfite, 2 mM H₂O₂, 200 µg/mL paraquat, and 2 M sorbitol. Additionally, growth under various carbon sources (12% maltose, 15% glucose + 15% fructose, 2% xylose, and 5% glycerol) and nitrogen sources (5 g/L isoleucine, proline, or histidine, with or without ammonium sulfate) was examined.

All conditions were tested at 20 °C, except in temperature-based assays, which were incubated at the corresponding temperatures. All stress media were prepared using minimal medium (MM; 20 g/L glucose, 0.67 g/L yeast nitrogen base with amino acids), supplemented with the appropriate stressor. pH conditions were adjusted using HCl or NaCl. For ethanol stress assays, ethanol was added post-autoclaving to final concentrations of 10%, 12%, or 15%. Nitrogen source variation was tested in MM lacking amino acids, supplemented with individual amino acids in the presence or absence of ammonium sulfate. For carbon source experiments, glucose was substituted with the respective alternative carbon sources in the MM formulation.

Yeast precultures were grown for two days at room temperature in MM containing 0.2% glucose (instead of 2%). For phenotypic screening, inocula were adjusted to an OD₆₀₀ of 0.1 in a final volume of 200 µL per well. OD_600_ was recorded every 24 hours over a 7-day period. Due to infrastructure limitations, only two time-point measurements were collected per day. Consequently, Max_Growth (i.e., maximum biomass production or maximum OD₆₀₀) was estimated using the `Growthcurver v0.3.1 R` package (Sprouffske and Wagner, 2016). We are aware that this approach does not capture growth rate, lag phase duration, or fermentation efficiency, parameters often more critical than final biomass in industrial applications (Barbosa *et al*., 2014). Future work should employ high-resolution growth curves and assess fermentation-relevant outputs (e.g., ethanol yield, metabolite profiles) to more fully evaluate industrial potential. Nevertheless, biomass production provides a valuable initial proxy for fitness and offers a practical starting point for identifying promising strains for further biotechnological development.

### Assessing mitotype effect

To assess whether mitochondrial inheritance affects growth fitness in allotetraploid strains, maximum growth values (Max_Growth) were analyzed using linear mixed-effects models (LMMs). All models included Experiment and Plate nested within Experiment as random effects to account for experimental batch and plate-to-plate variation. Analyses were restricted to allotetraploid (hybrid) strains only, excluding parental and control strains. Reference levels were defined as 20°C for condition, and mitotype inherited from *Saccharomyces* species.

We first compared the relative contribution of the nuclear and mitochondrial genomes to allotetraploid growth phenotypes, by using an additive LMM. Nuclear, mitochondrial genomes and conditions were included as fixed effects as follows: `Max_Growth ∼ NuclearPair + Mitotype * Condition + (1 | Experiment/Plate)`. For this model, the contributions of nuclear and mitochondrial components were compared using partial eta-squared (η²) effect sizes, using the function `eta_squared` from the `R` package `effectsize v1.0.0` (Ben-Shachar *et al*., 2020).

A second LMM was constructed with Max_Growth as the response variable, and Mitotype, Condition and their interactions as fixed effects: Mitotype * Condition `+ (1 | Experiment/Plate)`. Both LMMs were done using `R` packages `lme4 v1.1.35.5` (Bates *et al*., 2015) and `lmerTest v3.2.1` (Kuznetsova *et al*., 2017). In the second LMM, for each condition, an omnibus F-test of the Mitotype effect was obtained by evaluating this model within each level of Condition using the joint_tests function from `R` package `emmeans v1.10.6` (Lenth and Piaskowski, 2026). Given the size of the dataset (>12,000 observations), degrees of freedom for these tests were approximated using emmeans’ asymptotic (Wald) method; the global model’s overall ANOVA retained full Satterthwaite-approximated degrees of freedom via `lmerTest`. Resulting *p*-values were corrected for multiple testing across the 31 conditions using the Bonferroni method. For significant conditions, we also reported pairwise contrasts among the eight mitotypes within an allopolyploid, using the function ‘pairs’ from the `R` package `emmeans` with Tukey adjustment for multiple comparisons within each condition. For each condition, a mitotype was scored as a "win" against another mitotype when its estimated growth was significantly higher (p < 0.05, Tukey-adjusted), and a "loss" when significantly lower. A fitness score (wins minus losses) was calculated for each mitotype in each condition, and visualized as a heatmap using the `pheatmap v1.0.13` package (Kolde, 2025) in `R`, with rows (mitotypes) and columns (conditions) hierarchically clustered using Euclidean distance and complete linkage.

## Results

### Allotetraploids exhibit rapid sporulation and instability

To investigate the phenotypic impact of allopolyploidy in *Saccharomyces*, we employed the iterative hybrid production system (iHyPr) and mitoFIX molecular methods. These approaches allowed us to construct a comprehensive panel of allotetraploid yeasts, each involving distinct nuclear and mitochondrial genomic combinations (Figure 1, Supplementary Table 1).

Initially, we transformed diploid species using two differentially iHyPr plasmids, each carrying a drug-inducible *HO* endonuclease gene that facilitates efficient mating between diploids. Subsequently, these diploids were crossed and allotetraploids were selected based on the presence of both selectable drug markers (Figure 1). We constructed allotetraploids representing all combinations among eight *Saccharomyces* species, yielding up to 56 possible nuclear–mitochondrial species combinations and a total of 168 allotetraploids, with mostly three biological replicates generated for each combination. Due to limitations during transformation or recovery of *rho*^0^ strains, three categories of biological triplicates of allotetraploids were generated (Supplementary Table 1). The majority of allotetraploids (82.1%) comprises the intended experimental design, in which a parental *rho*^+^ species was crossed with three independently derived *rho*^0^ clones from another species. A second group (11.9%) comprises synthetic allotetraploids generated by reusing the same *rho*^0^ clones in one or two of the three independent crosses with the *rho*^+^ partner. The third group (6.0%) includes synthetic allotetraploids obtained by reusing existing synthetic allotetraploids, either twice (four allotetraploids) or three times (three allotetraploids), to complete the three required biological replicates. All allotetraploids and parental strains were validated using multiple complementary approaches.

Allotetraploids increased DNA content to the expected tetraploid (4x) state (Figure 2A). DNA content also correlated positively with cell size (Spearman rank-sum test R = 0.62, *p-*value < 2.2e^-16^; Supplementary Figure 3). However, the allotetraploid strains were highly unstable, and cells with reduced ploidy appeared within hours of growth in rich medium. This ploidy reduction likely occurred through sporulation, as supported by two lines of evidence. First, tetrads were observed microscopically, just growing in YPD plates at room temperature, indicating active sporulation (Figure 2C). Second, flow cytometry profiles revealed emerging cell populations with lower DNA content, consistent with the expected DNA content for a spore derived from a tetraploid cell (Figure 2C). These findings suggest that *Saccharomyces* allotetraploids are intrinsically unstable.

**Figure 2.**
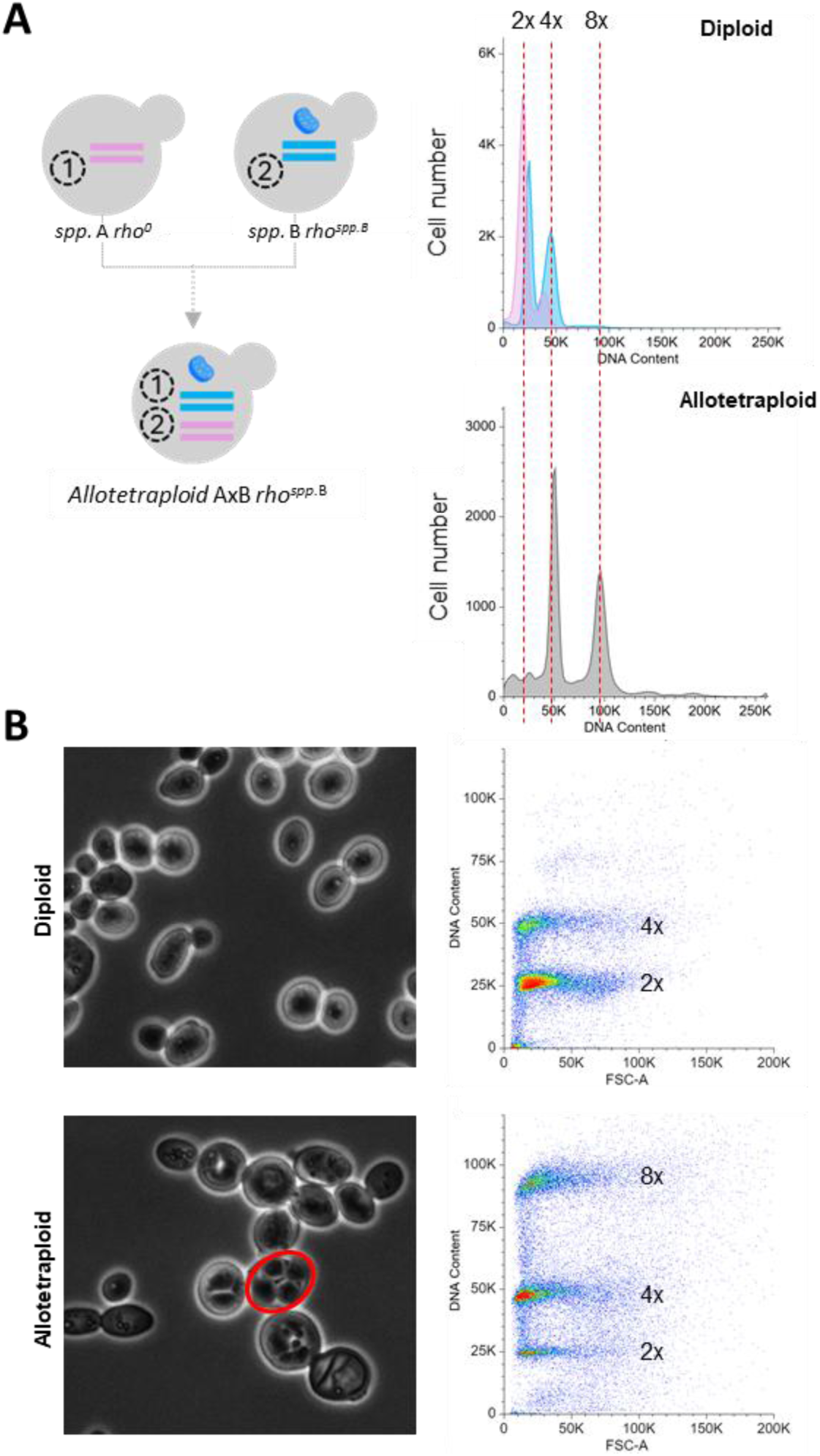
Highly unstable synthetic allotetraploids due to sporulation. **A**. The flow cytometry using SYTOX Green dye in both diploid parental strains and synthetic allotetraploid cells. The horizontal axis is the intensity of SYTOX Green fluorescence (proportional to DNA content). **B**. Light microscopy of yeast cells. A red circle indicates a tetrad. Quantification of DNA content by flow cytometry in both diploid and synthetic allotetraploid strains are displayed in different graphical representations in **A** and **B**. In **A**, the x-axis represents the fluorescence as a proxy of DNA content and in y-axis the number of cell counts. Each ploidy level is indicated above each pick. In **B**, the x-axis represents the forward scatter size (FSC-A), where an increased signal may indicate an increase in cell size or budding, and y-axis is the fluorescence as a proxy of DNA content. Diploid: *S. cerevisiae*, Allotetraploid: *S. cerevisiae* x *S. kudriavzevii*.

### Natural variation in industrially-relevant conditions reveal species-specific mitochondrial contributions to growth

To assess the contribution of each parent to allopolyploid phenotypes, we first compared the growth of wild-type *Saccharomyces* species across 31 diverse conditions, including different carbon and nitrogen sources, stressors, pH, and temperature (Supplementary Table 4). Optimal growth was observed in media with a pH between 3.5 and 7.5, 20°C, and in the presence of specific supplements such as proline or isoleucine (Figure 3A). Media supplemented with potassium metabisulfite, 3 mM or 7 mM FeCl₃ strongly inhibited growth across all tested strains (Figure 3A, Supplementary Table 5). Additionally, most species exhibited impaired growth at extreme temperatures (4 °C and 37 °C), under high ethanol concentrations, alkaline pH (e.g., pH 10), and in media containing sorbitol, glycerol, or carbon sources such as xylose or fructose (Figure 3A). Most strains failed to grow on maltose; however, *S. eubayanus* grew robustly, indicating the presence of functional maltose transporters and enzymes involved in maltose metabolism. Response to oxidative stress in the form of hydrogen peroxide was diverse among *Saccharomyces* species, with *S. jurei* displaying the greatest tolerance to oxidative stress induced by hydrogen peroxide (Figure 3A). In contrast, *S. eubayanus* and *S. uvarum* were the most resistant to oxidative stress caused by paraquat (a herbicide mimicking oxidative stress encountered during biomass propagation through the production of reactive oxygen species, ROS) (Figure 3A, Supplementary Table 4). Under elevated ethanol concentrations, *S. cerevisiae* and *S. paradoxus* exhibited the highest growth capacity. Notably, both *S. jurei* and *S. uvarum* showed improved growth in histidine-containing media without ammonium sulfate, whereas *S. jurei* performed best in histidine-containing media supplemented with ammonium sulfate.

**Figure 3.**
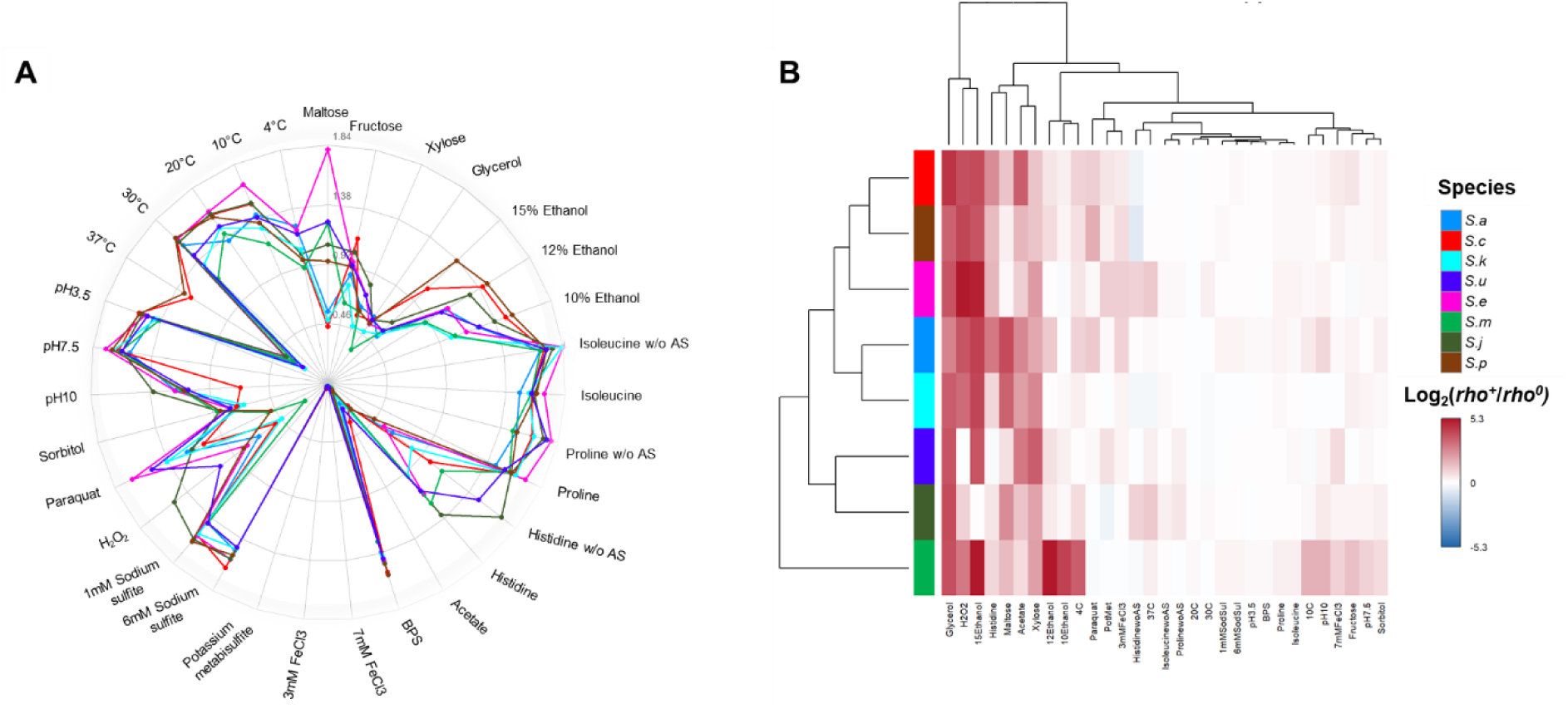
Diversity of wild parental phenotypes and the impact of mitochondrial DNA loss. **A.** Spider chart showing maximum biomass production values across different growth conditions for the diploid *Saccharomyces* parental strains (Supplementary Table 4). **B.** Heatmap showing log₂-transformed ratios of median maximum biomass production between *rho^+^* and *rho^0^* strains (log_2_[*rho^+^* / *rho^0^*] across *Saccharomyces* species and tested conditions, illustrating the effect of mitochondrial DNA removal. Rows represent *Saccharomyces* species and columns represent growth conditions. Species and conditions were clustered using hierarchical clustering with complete linkage on Euclidean distances. According to the color scale, red indicates higher growth in *rho^+^* compared to *rho^0^*; white indicates no change; and blue indicates higher growth in *rho^0^* relative to *rho^+^*. *S.a, S. arboricola; S.c, S. cerevisiae; S.k, S. kudriavzevii; S.u, S. uvarum; S.e, S. eubayanus; S.m, S. mikatae; S.j, S. jurei; S.p, S. paradoxus*.

To investigate the role of mitochondrial DNA in the growth of synthetic allotetraploids across different media, we first generated *rho^0^* derivatives of all eight diploid *Saccharomyces* species and performed an exploratory analysis of the effects of losing the mtDNA in the parental backgrounds. Comparison of *rho^+^* and *rho^0^* strains revealed that mitochondrial deficiency affected growth in several conditions, although the magnitude and specificity of these effects varied among genetic backgrounds (Figure 3B, Supplementary Figure 4). Most *rho*^0^ strains showed substantially reduced growth in glycerol, hydrogen peroxide, 15% ethanol, histidine, maltose, acetate and xylose. These phenotypes are consistent with the known inability of *rho^0^* cells to utilize non-fermentable carbon sources and with previously described roles of mitochondrial function in oxidative stress responses, ethanol tolerance, and redox balance during pentose utilization (Jin Yong-Su *et al*., 2004). Notably, *S. jurei* and *S. uvarum rho*^0^ strains displayed comparatively higher growth in the presence of hydrogen peroxide than other *Saccharomyces* species, indicating that loss of mtDNA did not abolish growth under oxidative stress in these backgrounds (Supplementary Figure 4, 5D). In contrast, *S. mikatae rho^0^* exhibited pronounced growth defects under additional conditions, including 10–15% ethanol, low temperatures (4–10°C), pH 10, and fructose, suggesting a particularly strong dependence on mitochondrial function across multiple environments (Figure 3B, Supplementary Figure 4, 5A-C). Overall, these results demonstrate that the contribution of mitochondrial DNA to growth and stress tolerance is highly species- and condition-dependent, and further validation will clarify the molecular mechanisms of tolerance in each species.

### Allopolyploidization retains parental traits and generates diverse phenotypic outcomes across environments

Following allopolyploidization using the previously phenotyped parental strains, the newly synthetic allotetraploids displayed extensive phenotypic diversification across a wide range of environmental and nutritional conditions (Figure 4C). Hierarchical clustering delineated two major groups of allotetraploids (maximum mean Silhouette width = 0.24 for K=2), largely driven by combinations containing *S. cerevisiae* (7/7) and *S. paradoxus* (5/7) parental genomes. The separation of these groups was primarily associated with the superior growth of these allotetraploids under high ethanol concentrations (10–15%) and elevated temperature (37°C), reflecting the contribution of the thermotolerant and ethanol-tolerant phenotypes of their parental species (Figure 3A, 4C).

**Figure 4.**
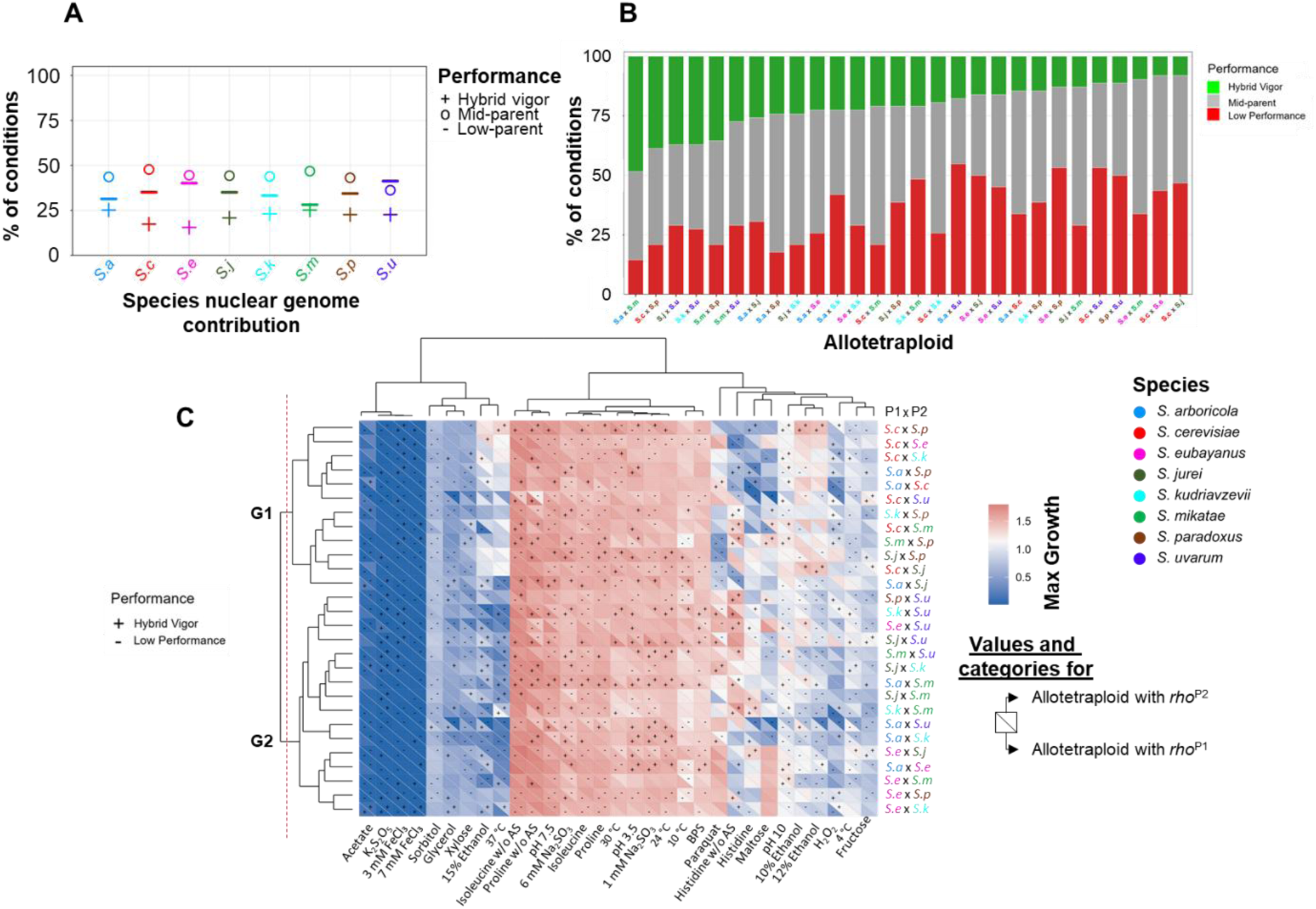
Genomic background drives growth variability in Saccharomyces *allotetraploids across conditions.* **A.** Dot plot showing the proportion of tested conditions assigned to each performance category: Mid-parent (median maximum growth intermediate between both parents, Supplementary Figure 6A), Hybrid Vigor (median maximum growth of allotetraploid exceeding that of the best parent, Supplementary Figure 6B), or Low Performance (median maximum growth lower than that of the worst parent). Allotetraploids were grouped according to the species contributing to the nuclear genome. **B.** Stacked bar plot depicting the proportion of tested conditions assigned to each performance category for each allotetraploid. **C.** Heatmap representing the median maximum growth values for each condition. The resulting allotetraploid x condition matrix was hierarchically clustered independently on both axes using Euclidean distance and complete-linkage agglomeration with `stats::hclust v4.4.1` package in `R v4.4`. For allotetraploid clustering analyses, the median maximum growth values obtained for the two reciprocal mitotype backgrounds were averaged, generating a single growth value for each nuclear genomic combination. For each allotetraploid combination, the lower-left triangle corresponds to the hybrid carrying the mitotype of parent 1 (P1), whereas the upper-right triangle corresponds to the reciprocal hybrid carrying the mitotype of parent 2 (P2). Symbols indicate categorical performance: “+” denotes hybrid vigor, and “-“ denotes low performance. The dendrogram was divided into two major groups after performing a mean Silhouette width test, which groups were delineated by a vertical dashed red line along the y-axis.

Most species combinations, excluding those involving *S. uvarum*, exhibited mid-parent heterosis (Figure 4A, Supplementary Figure 6A) in most of conditions (∼50%), defined as growth intermediate to that observed in both parents. Hybrid vigor in species combinations where usually lower of 25% of tested conditions (Figure 4A). However, there are particular conditions where we observed higher frequency of hybrid vigor (>35%) (Figure 4C): 6mM sodium sulfite, histidine without AS, 3mM FeCl3, K_2_S_2_O_5_, proline without AS, and pH 3.5. Despite hybrid vigor, certain stress conditions imposed a uniform constraint on growth across the entire collection. The strongest inhibitory effects were observed under acetate, potassium metabisulfite (K_2_S_2_O_5_), and iron chloride (3 and 7 mM FeCl_3_) (Figure 4C), conditions characterized by imposing oxidative and redox stress. The second group of inhibitory conditions were sorbitol, glycerol, xylose, 15% ethanol and 37°C, conditions that impose severe physiological stress and high energetic demands. These conditions challenge cellular homeostasis, respiration, redox balance, membrane integrity and protein stability, ultimately generating similar fitness outcomes. Even under these restrictive conditions, several allotetraploids maintained superior performance relative to their parents, indicating that allopolyploidy can partially buffer severe environmental stress.

Allotetraploids involving *S. arboricola* or *S. mikatae* parents usually generated hybrid vigor in 25% of conditions (Figure 4A, Supplementary Figure 6B). Allotetraploids generated by combining *S. arboricola* x *S. mikatae* presented hybrid vigor in ∼50% of conditions (Figure 4B), regardless of the mitotype (Figure 4C), indicating that this parental combination produces a particularly robust allopolyploid genotype across diverse environmental conditions. Other combinations, such as *S. cerevisiae* x *S. paradoxus*, *S. jurei* x *S. uvarum, S. kudriavzevii* x *S. uvarum* and *S. mikatae* x *S. paradoxus* were also quite vigorous in multiple conditions (Figure 4B-C). In contrast, allotetraploids containing *S. uvarum* were consistently associated with low-performance phenotypes across most conditions, suggesting pervasive antagonistic interactions between the *S. uvarum* genome and other parental backgrounds (Figure 4A-C). Notable exceptions were paraquat and histidine without AS conditions, where *S. uvarum*-containing allotetraploids outperformed most other hybrids. These phenotypes closely mirrored the superior performance of the parental *S. uvarum* strain under the same conditions (Figure 3A), indicating effective transmission of parental stress-response and nutrient-utilization traits to the allotetraploid background. A similar pattern was observed for *S. eubayanus*-containing allotetraploids under paraquat stress and maltose (Figure 4C), and for *S. mikatae*-containing allotetraploids in histidine medium lacking ammonium sulfate (Figure 4C), further supporting the notion that allotetraploids performance often reflects the ecological adaptations of specific parental species. Interestingly, the *S. mikatae* x *S. paradoxus* allotetraploid and *S. cerevisiae* x *S. mikatae* (*rho^Sm^*) also grew well on maltose (Figure 4C), suggesting some capacity of *S. mikatae* to consume maltose (Figure 3C).

Taken together, these results demonstrate that allotetraploid phenotypes emerge from a complex interplay of parental genomes, where both synergistic and antagonistic interactions shape fitness. Allopolyploid performance is highly condition-dependent, often recapitulating parental traits while also generating novel phenotypic combinations that are not predictable from either parent alone.

### Mitochondrial DNA inheritance impacts the growth of allotetraploids

Visual inspection of the growth profiles suggested that allotetraploids carrying alternative parental mitotypes exhibited distinct growth phenotypes for some conditions (Figure 4C). To disentangle the relative contributions of mitochondrial and nuclear genomes to allotetraploid phenotypes across environmental conditions, we applied LMMs. The additive model revealed significant effects of nuclear background (F_27,9632_ = 11.15, p < 2.2e-16), mitotype (F_7,9632_ = 22.72, P < 2.2e-16), environmental condition (F_30,9631_ = 906.55, P < 2.2e-16), and its interaction with mitotype (F_210,9632_ = 6.16, P < 2.2e-16). Environmental condition had the largest effect (partial η² = 0.74), followed by the interaction between mitotype and condition (partial η² = 0.12), whereas the nuclear genome (partial η² = 0.03) and mitotype (partial η² = 0.02) showed comparatively small main effects. Consistent with this context-dependent mitochondrial effect, mitotype (*rho*) significantly affected growth in 13 out of the 31 tested conditions (Supplementary Figure 7). These results indicate that mitochondrial inheritance plays an important role in shaping allotetraploid fitness, but that the magnitude of its effect depends strongly on environmental conditions.

Under ethanol stress, mitochondrial effects were particularly pronounced. Allotetraploids carrying *rho^Sc^* exhibited the highest growth, whereas those with *rho^Su^* consistently performed worse (Supplementary Figure 7). The effect of mitochondrial inheritance was also evident under oxidative stress. In the presence of hydrogen peroxide, allotetraploids carrying *rho^Sj^* exhibited the highest growth, whereas under paraquat stress, allotetraploids bearing *rho^Se^* consistently outperformed those carrying other mitotypes, highlighting a role for mitotype in modulating resistance to reactive oxygen species (Supplementary Figure 7). In histidine medium, allotetraploids carrying *rho^Sm^* and *rho^Sj^* generally performed better than other mitotypes both in the presence and absence of AS, whereas under AS-depleted conditions, those carrying *rho^Su^* showed the highest growth. Allotetraploids with *rho^Se^* performed best in maltose and proline without AS (Supplementary Figure 7). In low temperatures conditions, allotetraploids bearing *rho^Se^* or *rho^Sa^* performed best. In contrast, at higher temperatures *rho^Sc^* allotetraploids performed best. Mitotype-dependent differences were also observed in glycerol and xylose, where allotetraploids carrying *rho^Se^* and *rho^Sj^* exhibited the highest growth, respectively.

To further characterize these effects, we calculated a fitness score defined as the number of “wins” minus “losses” for each mitotype–condition combination (Figure 5). This analysis recapitulated patterns observed in the LMM (Supplementary Figure 7) and allowed direct comparison of mitotype performance across environments. The hierarchical clustering reflected the phylogenetic relationship of strains, except for *rho^Su^*.

**Figure 5.**
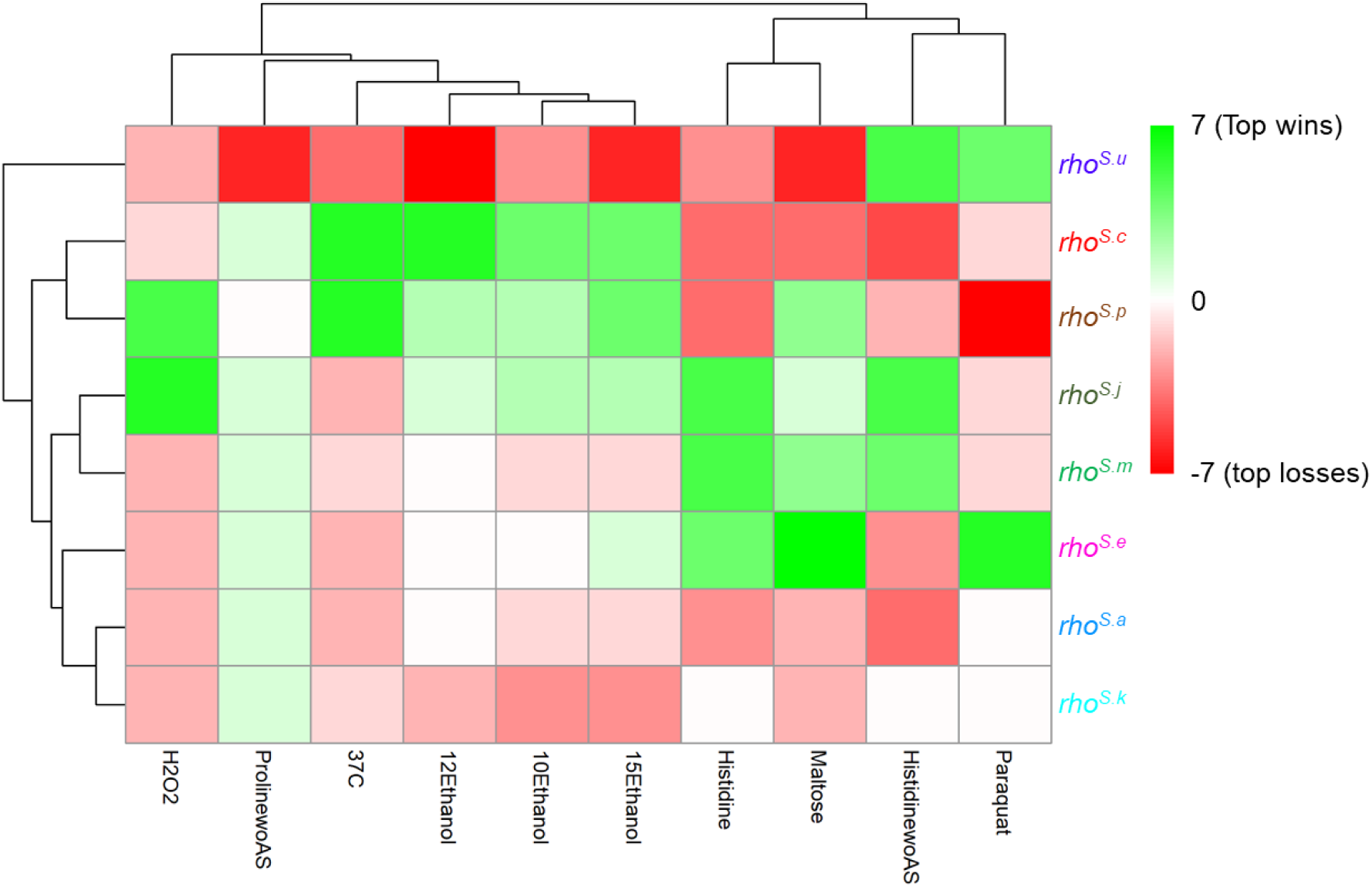
Condition-dependent growth as a function of mitotype inheritance. Heatmap shows fitness scores (number of wins minus losses) for each mitotype–condition combination for the 10 of 13 most significant conditions (Supplementary Figure 7). Green indicates higher relative fitness (growth advantage), red indicates lower relative fitness (growth disadvantage), and white represents neutral outcomes. Conditions are displayed along the x-axis, while mitotypes are shown on the y-axis. Independent Hierarchical clustering of both rows (mitotypes) and columns (conditions) was carried out using Euclidean distance and complete linkage. Maximum growth per condition for parents and allotetraploids, along with pairwise contrasts between mitotypes, is shown in Supplementary Figure 5.

Together, these results demonstrate that mitochondrial inheritance is a key, condition-dependent modulator of allotetraploid fitness. While environmental conditions remain the primary driver of growth, specific nuclear genome and especially mitotypes confer distinct advantages under particular environmental stresses, underscoring mitochondrial inheritance as a critical parameter that must be considered in the rational design of strains for biotechnological applications.

## Discussion

### Allopolyploids as a platform for strain improvement and diversity generation for the biotechnological industry

In this study, we constructed a large collection of allotetraploids combining the diploid genomes of all described *Saccharomyces* species (except for *S. chiloensis*). The successful construction of *Saccharomyces* allotetraploids with diverse nuclear and mitochondrial genomic combinations demonstrates the feasibility of using the iHyPr and mitoFIX molecular methods (Baker *et al*., 2019; Peris *et al*., 2020). Hybrid vigor in allotetraploids was present in ≤25% of all tested conditions, consistent with previous studies of interspecific yeast hybrids, where the phenotypic consequences of allopolyploidization are highly context-dependent and strongly influenced by the interaction between genomic background and environment (Peris et al., 2020, Stelkens *et al*., 2014; Bernardes *et al*., 2017; Brice *et al*., 2021; Haberkorn *et al*., 2026; Rinta-Harri *et al*., 2026). Allotetraploids exhibiting hybrid vigor may benefit from several genetic mechanisms, including positive epistatic interactions among divergent parental alleles, which can generate novel regulatory and metabolic states that enhances fitness as observed in plants, such as *Triticum aestivum* (Wu *et al*., 2025). Alternatively, heterosis may also arise through the complementation of deleterious alleles and the masking of recessive genetic load accumulated in different parental lineages, as reported in maize *Zea mays* (Sun *et al*., 2023). This highlight how the generation of synthetic allopolyploids is a potential tool for biotechnological applications, although is important to note that phenotyping was conducted in controlled laboratory media, though more efforts are needed to test these new strains in industrial settings, as these conditions present much complex stress combinations (Schwalbach *et al*., 2012) that may alter the relative importance of mitotype and reveal additional genotype-by-environment interactions not captured here.

Additionally, our allopolyploids demonstrated a high genome instability. This may be due to the presence of two divergent nuclear genomes, which can create imbalances in chromosome segregation, gene dosage, and regulatory networks, all of which can drive genomic instability (Otto, 2007). This behavior mirrors what has been observed in polyploid organisms across other kingdoms, where genome duplication often triggers genome downsizing, chromosomal loss, or diploidization as a means of restoring homeostasis (Wolfe, 2015). Moreover, carrying two copies of each parental genome might impose an energetic cost triggering stress responses (Barker *et al*., 2025) which is likely the reason why sporulation is observed in all allotetraploid combinations tending to restore diploid levels, while overcoming infertility observed in allodiploids (Naseeb *et al*., 2021). However, the rapid sporulation and genomic instability observed in these allotetraploids represent a double-edge sword. On one side, the genomic instability creates opportunities for innovation, as sporulation generates extensive diversity that can be leveraged in breeding and adaptive evolution programs to identify strains with improved industrial performance (Krogerus, Seppänen-Laakso, *et al*., 2017) that after some generations can improve the growth kinetics compared to parental strains (Peris, Moriarty, *et al*., 2017; Peris *et al*., 2020; Molinet *et al*., 2024) or provide novel metabolic states, stress responses, and fermentation-related traits such as aroma and flavour compound production (Bellon *et al*., 2013). This process may act as a natural filter, favoring genomic configurations that are more compatible or robust expanding the phenotypic landscape for strain improvement (Gupte *et al*., 2023). On the other side, they pose a major challenge for industrial deployment, where genetic stability over repeated propagation cycles is essential for maintaining process robustness and consistent expression of desirable industrially-relevant traits (Gorter de Vries *et al*., 2019). This limitation may require further strain stabilization through adaptive evolution or the isolation of non-sporulating derivates. Additionally, this level of sporulation activity represents a barrier to generate initial higher ploidy levels. In our previous six-species allopolyploid designs, where we pursued allododecaploid strains, we only reached the alloctaploid state (Peris *et al*., 2020). Similarly, other studies have reported lower ploidy levels than originally anticipated following polyploid construction (Rinta-Harri *et al*., 2026). These observations suggest that sporulation and ploidy reduction constitute pervasive mechanisms that constrain the maintenance of highly polyploid genomes in synthetic *Saccharomyces* polyploids.

Nonetheless, sporulation enables classical genetic approaches such as QTL mapping, facilitating the identification of genes underlying complex traits (Naseeb *et al*., 2021). In *S. cerevisiae*, meiotic progeny derived from crosses have been extensively used to map Quantitative Trait Loci (QTLs) controlling industrially relevant phenotypes (Salinas *et al*., 2012). Thus, while genomic instability may limit long-term tetraploid maintenance, the ability to sporulate provides a key advantage for generating diversity and dissecting genotype–phenotype relationships, making allotetraploids highly valuable tools in yeast biotechnology.

### The S.H.E.L. platform: select, hybridize, evolve and learn cycle

Although synthetic allotetraploids or higher ploidy hybrids will help dissecting genotype–phenotype relationships, engineering complex traits such as ethanol or temperature tolerance are complex (Voordeckers *et al*., 2015; Weiss *et al*., 2018). Biotechnological strain development often focuses on optimizing a single genotype for multiple conditions or traits, an approach that is inherently limited by the genetic potential of that individual strain. Hybridization allows rapid exploration of the phenotype space, combining traits of multiples strains, without requiring genetic engineering. Several of the species included in this study have already attracted considerable interest for industrial applications, either individually or as components of natural and synthetic interspecies hybrids (Magalhães *et al*., 2017; Drężek *et al*., 2024; Zavaleta Vasni *et al*., 2024). In particular, *S. cerevisiae, S. uvarum, S. kudriavzevii,* and *S. eubayanus* are recurrent components of industrial hybrids isolated from brewing, wine, and other fermentation environments, highlighting their potential as reservoirs of desirable traits (Peris *et al*., 2018; Langdon *et al*., 2019). The rational behind our species selection was therefore to exploit the phenotypes naturally present across the genus *Saccharomyces* (Paliwal *et al*., 2014; Peter *et al*., 2018; Peris *et al*., 2023). For example, *S. cerevisiae* contributes high ethanol and thermotolerance, traits that are highly desirable for alcoholic fermentations and biofuel production (Lairón-Peris *et al*., 2020). *S. eubayanus* is renowned for its ability to metabolize maltose (Quintrel *et al*., 2025) and to grow at low temperatures, making it particularly valuable for brewing applications (Baker *et al*., 2019). In contrast, *S. uvarum* and *S. kudriavzevii* are associated with enhanced glycerol production and distinct aromatic profiles that are attractive for wine fermentations (Peris *et al*., 2016; Minebois *et al*., 2020). Our results further suggest that less-studied species may also represent valuable sources of industrially relevant variation, as traits such as oxidative stress tolerance, growth under nitrogen limitation, and cold adaptation were particularly enriched in *S. jurei*, *S. mikatae*, and *S. arboricola* backgrounds.

Careful strain selection is essential, as the optimal choice depends on the intended biotechnological application. For example, our results show that allotetraploids generated with our selected *S. uvarum* strain performed poorly across most conditions, yet in histidine medium without AS they exhibited the highest growth of all genotypes. This observation highlights the limitations of using a single representative strain to generalizability of findings to the species. We envision that, as an increasing number of yeast strains are phenotypically and genomically characterized, they can be incorporated into iterative applications of a framework we term the SHEL cycle (Select–Hybridize–Evolve–Learn), conceptually analogous to the Design–Build–Test–Learn cycle widely used in synthetic biology. However, rather than focusing on direct genetic engineering, the SHEL cycle is centered on iterative hybridization and evolutionary improvement. This framework integrates the selection of diverse parental strains, the generation of synthetic hybrids and allopolyploids, experimental evolution to enhance performance and stabilize genomes, and the systematic learning of genotype–phenotype relationships. Importantly, the SHEL cycle explicitly considers both nuclear and extrachromosomal genomes (i.e. mtDNA), as key variables in the rational design of new synthetic crosses for biotechnological applications.

### The underestimated mitochondrial inheritance in synthetic allopolyploids and its biotechnological consequences

Extrachromosomal genomes, particularly mitochondrial DNA should also be considered. Despite their well-established role in shaping phenotypic variation, mitochondrial genomes have been largely overlooked in most recent studies on *Saccharomyces* allopolyploids and hybrids (Gyurchev *et al*., 2022; Haberkorn *et al*., 2026; Martínez and Lang, 2026; Rinta-Harri *et al*., 2026). Mitochondrial variation is not only important in interspecific crosses, but can also play an important role in crosses between genetically divergent strains within species, where distinct mitotypes can generate phenotypic differences (Paliwal *et al*., 2014). We demonstrated that mitotype influenced the growth in 42% of conditions. Mitotype-associated traits previously described in *Saccharomyces*, including maltose utilization (Molinet *et al*., 2024) and low-temperature tolerance in *S. eubayanus* (Baker *et al*., 2019), and high temperature tolerance in *S. cerevisiae* (Baker *et al*., 2019; Li *et al*., 2019) were consistently recapitulated in our study across a wide range of allotetraploid backgrounds rather than being restricted to a few species combinations. Unexpectedly, the *S. uvarum* mitotype did not confer superior growth at low temperature, despite previous reports linking *S. uvarum* mitochondria to cold tolerance (Li *et al*., 2019). This suggests that the particular *S. uvarum* strain used here may not be representative of the species’ typical mitochondrial phenotype, as discussed above. Specific molecular mismatches, such as the incompatibility between the mitochondrial and nuclear-encoded proteins (Jhuang *et al*., 2017) or failures in mitonuclear coordination (Lee *et al*., 2008) might contribute to the dysfunction and reduced growth of *S. uvarum* allotetraploids.

Our approach also identified mitochondrial genomes from less-studied *Saccharomyces* species with considerable biotechnological potential. In particular, mitotypes from *S. jurei*, *S. arboricola*, and *S. mikatae* conferred advantages under oxidative stress, respiratory growth conditions, alternative carbon sources such as xylose, and nitrogen-limited environments. Elucidating the molecular mechanisms underlying these mitotype-specific effects represents a promising avenue for future research and may uncover novel targets for strain improvement.

## Conclusions

This work shows that adaptability in *Saccharomyces* arises from a multi-layered interaction between nuclear and mitochondrial genomes, strongly shaped by environmental context. While environment explains the largest part of the phenotypic variation observed in allotetraploids, mitochondrial variation consistently emerges as an additional and condition-dependent driver of performance, enhancing or constraining key industrial traits. Importantly, these results shift the emphasis from optimizing individual genotypes to harnessing natural nuclear and mitochondial genomic diversity as the primary source of functional innovation: allopolyploid systems should be understood as tools to amplify and recombine existing natural variation across both nuclear and mitochondrial genomes. In future SHEL cycle frameworks, the strain development should explicitly consider mitotype choice to unlock optimal combinations for specific industrial applications such as fermentation efficiency, stress tolerance, and substrate utilization.

## Data availability

Raw sequencing, flow cytometry, and spectrometry data have been deposited in Figshare doi: 10.6084/m9.figshare.30598748. HyPr plasmids are deposited in Addgene, deposit number 77444.

## Acknowledgements

We thank Rafal Ciosk (Department of Biosciences, University of Oslo) for providing access to the spectrophotometer, and Yan Zhang (Department of Pathology, University of Oslo) for facilitating flow cytometry analyses. We are grateful to Cecilie Mathiesen, Maria Chiara Di Luca (Department of Biosciences, University of Oslo) for technical assistance; Inger Skrede and Håvard Kauserud (Department of Biosciences, University of Oslo) for departmental support. We also thank Héctor García Martin (Lawrence Berkeley National Laboratory and Director of Data Science and Modeling at the Joint BioEnergy Institute, USA) for valuable discussions that contributed to the development of the SHEL cycle concept. We also acknowledge the valuable feedback received during project development from members of the Oslo Mycology Group and FunGIALab at University of Oslo and Institute of Agrochemistry and Food Technology, respectively. This work was supported by the Research Council of Norway (grant RCN 324253, project PloidYeast) and *Conselleria de Cultura, Educación y Ciencia, Generalitat Valenciana* CIDEGENT/2021/039 & CIESGT/2024/012. S.O.M. was funded by grant RCN 324253. D.P. was supported by multiple grants, including: Spanish government MCIN/AEI to the Center of Excellence Accreditation Severo Ochoa (CEX2021-001189-S-20-10); TED2021-131349B-I00 (*Proyectos Estratégicos Orientados a la Transición Ecológica y a la Transición Digital*), funded by MCIN/AEI/10.13039/501100011033 and by the European Union NextGenerationEU/PRTR; Spanish Ministry of Science, Innovation and Universities (MICIU/AEI/10.13039/501100011033) through grant PID2025-169211OB-I00. DP receives additional funding from CSIC COOPA24016, FUNDS2024023 and AGAIN25019. R.S. and C.H. were supported by a Swedish Research Council Project Grant (2022-03427) and a Knut and Alice Wallenberg Foundation Grant (2024.0216).

## Author contributions

S.O.M. conducted the experimental work, including parental strain engineering, generation of synthetic allotetraploid strains, and strain validation (with minor contributions from B.C.); S.O.M. generated the data. S.O.M. and D.P. performed the growth curve analyses; C.H., R.S., and D.P. carried out the statistical analyses using linear mixed models (LMM) analyses; S.O.M. and D.P. jointly conceived and designed the study. D.P. supervised the project. S.O.M. and D.P. wrote the manuscript, with editorial input from R.S. and C.H. All authors reviewed and approved the final version of the manuscript.

## Competing interests

The Wisconsin Alumni Research Foundation holds patents titled “Synthetic yeast cells and methods of making and using same” (describing the HyPr and iHyPr methods, with D.P. as inventor; US patent No. US20180127784A1, valid until November 7, 2037) and “Yeast strains with selected or altered mitotypes and methods of making and using the same” (describing the mitoFIX method, also with D.P. as inventor; US patent No. US20200048645A1, valid until October 7, 2039). The remaining authors declare no competing interests.

The authors used LLMs to assist with language editing and grammar improvement. All content was reviewed, edited, and approved by the authors, who take full responsibility for the final text.

## Additional information

**Supplementary information** is available for this paper at XXX.

**Correspondence** and requests for materials should be addressed to D.P.

## Synthetic allotetraploids Supporting information captions

***Supplementary Figure 1.*** *Phenotype of the rho^0^ yeast strains*.

***Supplementary Figure 2.*** *Confirmation of newly synthetized allotetraploid strains*

***Supplementary Figure 3.*** *Positive correlation between DNA content (GFP-A) and cell size (FCS-A)*.

***Supplementary Figure 4.*** *Differential impact of mitochondrial DNA loss on growth across* Saccharomyces *species*.

***Supplementary Figure 5.*** *Environmental conditions revealing mitotype-dependent phenotypic variation in synthetic allotetraploids*.

***Supplementary Figure 6.*** *Best parent heterosis is limited among environmental conditions*.

***Supplementary Figure 7.*** *Mitochondrial DNA origin has an impact on allotetraploid growth across environments*.

***Supplementary Table 1.*** *Strains used in this work.*

***Supplementary Table 2.*** *Plasmids used in this work.*

***Supplementary Table 3.*** *Oligonucleotides used in this work.*

***Supplementary Table 4.*** *Tested media*.

***Supplementary Table 5.*** *Summary statistics of the inferred maximum biomass (Max_Growth)*.

## Notes

https://doi.org/10.6084/m9.figshare.30598748

## References

Adams, K. (2003) Evolution of mitochondrial gene content: gene loss and transfer to the nucleus. Mol Phylogenet Evol 29: 380–395.

Alexander, W.G., Peris, D., Pfannenstiel, B.T., Opulente, D.A., Kuang, M., and Hittinger, C.T. (2016) Efficient engineering of marker-free synthetic allotetraploids of Saccharomyces. Fungal Genet Biol 89: 10–17.

Bachinskaya (1914) Saccharomyces paradoxus Bach.-Raich., 1914. 1: 231.

Baker, E.P., Peris, D., Moriarty, R.V., Li, X.C., Fay, J.C., and Hittinger, C.T. (2019) Mitochondrial DNA and temperature tolerance in lager yeasts. Sci Adv 5: eaav1869.

Barbosa, C., Lage, P., Vilela, A., Mendes-Faia, A., and Mendes-Ferreira, A. (2014) Phenotypic and metabolic traits of commercial Saccharomyces cerevisiae yeasts. AMB Express 4: 39.

Barker, J., Murray, A., and Bell, S.P. (2025) Cell integrity limits ploidy in budding yeast. G3 GenesGenomesGenetics 15: jkae286.

Bates, D., Mächler, M., Bolker, B., and Walker, S. (2015) Fitting Linear Mixed-Effects Models Using **lme4**. J Stat Softw 67:.

Beijerinck, M.W. (1898) Saccharomyces uvarum Beijerinck.

Bellon, J.R., Schmid, F., Capone, D.L., Dunn, B.L., and Chambers, P.J. (2013) Introducing a new breed of wine yeast: interspecific hybridisation between a commercial *Saccharomyces cerevisiae* wine yeast and *Saccharomyces mikatae*. PLoS ONE 8: e62053.

Bendixsen, D.P., Peris, D., and Stelkens, R. (2021) Patterns of genomic instability in interspecific yeast hybrids with diverse ancestries. Front Fungal Biol 2: 52.

Ben-Shachar, M.S., Lüdecke, D., and Makowski, D. (2020) effectsize: Estimation of Effect Size Indices and Standardized Parameters. J Open Source Softw 5: 2815.

Bernardes, J.P., Stelkens, R.B., and Greig, D. (2017) Heterosis in hybrids within and between yeast species. J Evol Biol 30: 538–548.

Brice, C., Zhang, Z., Bendixsen, D., and Stelkens, R. (2021) Hybridization outcomes have strong genomic and environmental contingencies. Am Nat.

Burton, R.S. and Barreto, F.S. (2012) A disproportionate role for mt DNA in D obzhansky–M uller incompatibilities? Mol Ecol 21: 4942–4957.

Chen, Z.J. (2007) Genetic and Epigenetic Mechanisms for Gene Expression and Phenotypic Variation in Plant Polyploids. Annu Rev Plant Biol 58: 377–406.

Comai, L. (2005) The advantages and disadvantages of being polyploid. Nat Rev Genet 6: 836–846.

Dandage, R., Berger, C.M., Gagnon-Arsenault, I., Moon, K.-M., Stacey, R.G., Foster, L.J., and Landry, C.R. (2021) Frequent Assembly of Chimeric Complexes in the Protein Interaction Network of an Interspecies Yeast Hybrid. Mol Biol Evol 38: 1384–1401.

Desmazières, J.B. (1827) Recherches microscopiques et physiologiques sur le genre *Mycoderma*. Ann Sci Nat 10: 42–67.

Drężek, K., Antunovics, Z., and Grabiec, A.K. (2024) Novel Saccharomyces uvarum x Saccharomyces kudriavzevii synthetic hybrid with enhanced 2-phenylethanol production. Microb Cell Factories 23: 203.

Eckert-Boulet, N., Rothstein, R., and Lisby, M. (2011) Cell Biology of Homologous Recombination in Yeast. In DNA Recombination. Methods in Molecular Biology. Tsubouchi, H. (ed). Totowa, NJ: Humana Press, pp. 523–536.

FlowJo (1996).

Gallone, B., Steensels, J., Mertens, S., Dzialo, M.C., Gordon, J.L., Wauters, R., et al. (2019) Interspecific hybridization facilitates niche adaptation in beer yeast. Nat Ecol Evol 3: 1562–1575.

Gorter de Vries, A.R., Voskamp, M.A., van Aalst, A.C.A., Kristensen, L.H., Jansen, L., van den Broek, M., et al. (2019) Laboratory evolution of a *Saccharomyces cerevisiae* x *S. eubayanus* hybrid under simulated Lager-brewing conditions. Front Genet 10: 242.

Gupte, A.P., Pierantoni, D.C., Conti, A., Donati, L., Basaglia, M., Casella, S., et al. (2023) Renewing Lost Genetic Variability with a Classical Yeast Genetics Approach. J Fungi 9: 264.

Gyurchev, N.Y., Coral-Medina, Á., Weening, S.M., Almayouf, S., Kuijpers, N.G.A., Nevoigt, E., and Louis, E.J. (2022) Beyond *Saccharomyces pastorianus* for modern lager brews: Exploring non-*cerevisiae Saccharomyces* hybrids with heterotic maltotriose consumption and novel aroma profile. Front Microbiol 13:.

Haase, S.B. and Reed, S.I. (2002) Improved flow cytometric analysis of the budding yeast cell cycle. Cell Cycle Georget Tex 1: 132–136.

Haberkorn, C., Gettle, N., Elsen, J., Medina Chavez, N.O., Sivigny, J., Baselga-Cervera, B., et al. (2026) Adaptive benefits of hybridization in Saccharomyces yeast are constrained by genomic background and depend on temperature. Evolution qpag108.

Hewitt, S.K., Donaldson, I.J., Lovell, S.C., and Delneri, D. (2014) Sequencing and Characterisation of Rearrangements in Three S. pastorianus Strains Reveals the Presence of Chimeric Genes and Gives Evidence of Breakpoint Reuse. PLoS ONE 9: e92203.

Jhuang, H., Lee, H., and Leu, J. (2017) Mitochondrial–nuclear co-evolution leads to hybrid incompatibility through pentatricopeptide repeat proteins. EMBO Rep 18: 87–101.

Jin Yong-Su, Laplaza Jose M., and Jeffries Thomas W. (2004) Saccharomyces cerevisiae Engineered for Xylose Metabolism Exhibits a Respiratory Response. Appl Environ Microbiol 70: 6816–6825.

Kolde, R. (2025) pheatmap: Pretty Heatmaps.

Krogerus, K., Magalhães, F., Vidgren, V., and Gibson, B. (2015) New lager yeast strains generated by interspecific hybridization. J Ind Microbiol Biotechnol 42: 769–778.

Krogerus, K., Magalhães, F., Vidgren, V., and Gibson, B. (2017) Novel brewing yeast hybrids: creation and application. Appl Microbiol Biotechnol 101: 65–78.

Krogerus, K., Seppänen-Laakso, T., Castillo, S., and Gibson, B. (2017) Inheritance of brewing-relevant phenotypes in constructed *Saccharomyces cerevisiae* x *Saccharomyces eubayanus* hybrids. Microb Cell Factories 16: 66.

Kuznetsova, A., Brockhoff, P.B., and Christensen, R.H.B. (2017) **lmerTest** Package: Tests in Linear Mixed Effects Models. J Stat Softw 82:.

Lairón-Peris, M., Pérez-Través, L., Muñiz-Calvo, S., Guillamón, J.M., Heras, J.M., Barrio, E., and Querol, A. (2020) Differential contribution of the parental genomes to a *S. cerevisiae* x *S. uvarum* hybrid, inferred by phenomic, genomic, and transcriptomic analyses, at different industrial stress conditions. Front Bioeng Biotechnol 8: 129.

Langdon, Q.K., Peris, D., Baker, E.P., Opulente, D.A., Nguyen, H.-V., Bond, U., et al. (2019) Fermentation innovation through complex hybridization of wild and domesticated yeasts. Nat Ecol Evol 3: 1576–1586.

Leducq, J.-B., Charron, G., Diss, G., Gagnon-Arsenault, I., Dubé, A.K., and Landry, C.R. (2012) Evidence for the Robustness of Protein Complexes to Inter-Species Hybridization. PLoS Genet 8: e1003161.

Leducq, J.-B., Henault, M., Charron, G., Nielly-Thibault, L., Terrat, Y., Fiumera, H.L., et al. (2017) Mitochondrial recombination and introgression during speciation by hybridization. Mol Biol Evol 34: 1947–1959.

Lee, H.-Y., Chou, J.-Y., Cheong, L., Chang, N.-H., Yang, S.-Y., and Leu, J.-Y. (2008) Incompatibility of Nuclear and Mitochondrial Genomes Causes Hybrid Sterility between Two Yeast Species. Cell 135: 1065–1073.

Lenth, R.V. and Piaskowski, J. (2026) emmeans: Estimated Marginal Means, aka Least-Squares Means.

Li, X.C., Peris, D., Hittinger, C.T., Sia, E.A., and Fay, J.C. (2019) Mitochondria-encoded genes contribute to the evolution of heat and cold tolerance among *Saccharomyces* species. Sci Adv 5: eaav1848.

Libkind, D., Hittinger, C.T., Valério, E., Gonçalves, C., Dover, J., Johnston, M., et al. (2011) Microbe domestication and the identification of the wild genetic stock of lager-brewing yeast. Proc Natl Acad Sci 108: 14539–14544.

Libkind, D., Peris, D., Cubillos, F.A., Steenwyk, J.L., Opulente, D.A., Langdon, Q.K., et al. (2020) Into the wild: new yeast genomes from natural environments and new tools for their analysis. FEMS Yeast Res 20: foaa008.

Magalhães, F., Krogerus, K., Vidgren, V., Sandell, M., and Gibson, B. (2017) Improved cider fermentation performance and quality with newly generated *Saccharomyces cerevisiae* x *Saccharomyces eubayanus* hybrids. J Ind Microbiol Biotechnol 1–11.

Malina, C., Larsson, C., and Nielsen, J. (2018) Yeast mitochondria: an overview of mitochondrial biology and the potential of mitochondrial systems biology. FEMS Yeast Res 18: foy040.

Martínez, A.A. and Lang, G.I. (2026) Weak pervasive incompatibilities and compensatory adaptation drive hybrid genome evolution in yeast. bioRxiv 2026.01.07.698244.

Minebois, R., Pérez-Torrado, R., and Querol, A. (2020) A time course metabolism comparison among *Saccharomyces cerevisiae*, S. uvarum and S. kudriavzevii species in wine fermentation. Food Microbiol 90: 103484.

Molinet, J., Navarrete, J.P., Villarroel, C.A., Villarreal, P., Sandoval, F.I., Nespolo, R.F., et al. (2024) Wild Patagonian yeast improve the evolutionary potential of novel interspecific hybrid strains for lager brewing. Plos Genet 20: e1011154.

Naseeb, S., Alsammar, H., Burgis, T., Donaldson, I., Knyazev, N., Knight, C., and Delneri, D. (2018) Whole Genome Sequencing, *de Novo* Assembly and Phenotypic Profiling for the New Budding Yeast Species *Saccharomyces jurei*. G3 GenesGenomesGenetics 8: 2967–2977.

Naseeb, S., James, S.A., Alsammar, H., Michaels, C.J., Gini, B., Nueno-Palop, C., et al. (2017) Saccharomyces jurei sp. nov., isolation and genetic identification of a novel yeast species from Quercus robur. Int J Syst Evol Microbiol 67: 2046–2052.

Naseeb, S., Visinoni, F., Hu, Y., Hinks Roberts, A.J., Maslowska, A., Walsh, T., et al. (2021) Restoring fertility in yeast hybrids: Breeding and quantitative genetics of beneficial traits. Proc Natl Acad Sci 118: e2101242118.

Naumov, G.I., James, S.A., Naumova, E.S., Louis, E.J., and Roberts, I.N. (2000) Three new species in the *Saccharomyces sensu stricto* complex: *Saccharomyces cariocanus*, *Saccharomyces kudriavzevii* and *Saccharomyces mikatae*. Int J Syst Evol Microbiol 50: 1931–1942.

Otto, S.P. (2007) The Evolutionary Consequences of Polyploidy. Cell 131: 452–462.

Paliwal, S., Fiumera, A.C., and Fiumera, H.L. (2014) Mitochondrial-nuclear epistasis contributes to phenotypic variation and coadaptation in natural isolates of *Saccharomyces cerevisiae*. Genetics 198: 1251–1265.

Peña, T.A., Villarreal, P., Agier, N., De Chiara, M., Barría, T., Urbina, K., et al. (2024) An integrative taxonomy approach reveals Saccharomyces chiloensis sp. nov. as a newly discovered species from Coastal Patagonia. PLOS Genet 20: e1011396.

Peris, D., Alexander, W.G., Fisher, K.J., Moriarty, R.V., Basuino, M.G., Ubbelohde, E.J., et al. (2020) Synthetic hybrids of six yeast species. Nat Commun 11: 2085.

Peris, D., Arias, A., Orlić, S., Belloch, C., Pérez-Través, L., Querol, A., and Barrio, E. (2017) Mitochondrial introgression suggests extensive ancestral hybridization events among Saccharomyces species. Mol Phylogenet Evol 108: 49–60.

Peris, D., Belloch, C., Lopandić, K., Álvarez-Pérez, J.M., Querol, A., and Barrio, E. (2012) The molecular characterization of new types of *Saccharomyces cerevisiae* × *S. kudriavzevii* hybrid yeasts unveils a high genetic diversity. Yeast 29: 81–91.

Peris, D., Lopes, C.A., Belloch, C., Querol, A., and Barrio, E. (2012) Comparative genomics among Saccharomyces cerevisiae × Saccharomyces kudriavzevii natural hybrid strains isolated from wine and beer reveals different origins. BMC Genomics 13: 407.

Peris, D., Moriarty, R.V., Alexander, W.G., Sylvester, K., Sardi, M., Libkind, D., et al. (2017) Hybridization and adaptive evolution of diverse *Saccharomyces* species for cellulosic biofuel production. Biotechnol Biofuels 10: 78.

Peris, D., Pérez-Torrado, R., Hittinger, C., Barrio, E., and Querol, A. (2018) On the origins and industrial applications of *Saccharomyces cerevisiae* x *Saccharomyces kudriavzevii* hybrids. Yeast 5: 51–69.

Peris, D., Pérez-Través, L., Belloch, C., and Querol, A. (2016) Enological characterization of Spanish *Saccharomyces kudriavzevii* strains, one of the closest relatives to parental strains of winemaking and brewing *S. cerevisiae* x *S. kudriavzevii* hybrids. Food Microbiol 53: 31–40.

Peris, D., Sylvester, K., Libkind, D., Gonçalves, P., Sampaio, J.P., Alexander, W.G., and Hittinger, C.T. (2014) Population structure and reticulate evolution of *Saccharomyces eubayanus* and its lager-brewing hybrids. Mol Ecol 23: 2031–2045.

Peris, D., Ubbelohde, E.J., Kuang, M.C., Kominek, J., Langdon, Q.K., Adams, M., et al. (2023) Macroevolutionary diversity of traits and genomes in the model yeast genus Saccharomyces. Nat Commun 14: 690.

Peter, J., De Chiara, M., Friedrich, A., Yue, J.-X., Pflieger, D., Bergström, A., et al. (2018) Genome evolution across 1,011 Saccharomyces cerevisiae isolates. Nature 556: 339–344.

Piatkowska, E.M., Naseeb, S., Knight, D., and Delneri, D. (2013) Chimeric Protein Complexes in Hybrid Species Generate Novel Phenotypes. PLoS Genet 9: e1003836.

Piccinini, G., Iannello, M., Puccio, G., Plazzi, F., Havird, J.C., and Ghiselli, F. (2021) Mitonuclear Coevolution, but not Nuclear Compensation, Drives Evolution of OXPHOS Complexes in Bivalves. Mol Biol Evol 38: 2597–2614.

Quintrel, P., Muñoz-Guzmán, F., Villarreal, P., Peña, T.A., Garate, N.I., Muñoz-Tapia, C., et al. (2025) Allelic variation in MAL33 drives ecological adaptation of maltose metabolism in Saccharomyces eubayanus. bioRxiv 2025.09.15.676268.

R Development Core Team (2010) R: a language and environment for statistical computing. Httpwww R-Proj Org.

Ramírez-Aroca, L. (2019) Influencia del genoma mitocondrial en el fenotipo de híbridos interespecíficos durante condiciones industriales.

Rinta-Harri, K., Koponen, T., Mojzita, D., Jouhten, P., Liti, G., and Krogerus, K. (2026) Influence of ploidy and genetic background on stress tolerance of intraspecific yeast hybrids. Microb Biotechnol 19: e70337.

Salinas, F., Cubillos, F.A., Soto, D., Garcia, V., Bergström, A., Warringer, J., et al. (2012) The Genetic Basis of Natural Variation in Oenological Traits in Saccharomyces cerevisiae. PLoS ONE 7: e49640.

Schwalbach, M.S., Keating, D.H., Tremaine, M., Marner, W.D., Zhang, Y., Bothfeld, W., et al. (2012) Complex physiology and compound stress responses during fermentation of Alkali-Pretreated Corn Stover Hydrolysate by an *Escherichia coli* ethanologen. Appl Environ Microbiol 78: 3442–3457.

Shen, X.-X., Steenwyk, J.L., LaBella, A.L., Opulente, D.A., Zhou, X., Kominek, J., et al. (2020) Genome-scale phylogeny and contrasting modes of genome evolution in the fungal phylum Ascomycota. Sci Adv 6: eabd0079.

Sherman, F. (1963) Respiration-deficient mutants of yeast. Genetics 48: 375–385.

Sickmann, A., Reinders, J., Wagner, Y., Joppich, C., Zahedi, R., Meyer, H.E., et al. (2003) The proteome of *Saccharomyces cerevisiae* mitochondria. Proc Natl Acad Sci 100: 13207–13212.

Soltis, P.S. and Soltis, D.E. (2009) The Role of Hybridization in Plant Speciation. Annu Rev Plant Biol 60: 561–588.

Sprouffske, K. and Wagner, A. (2016) Growthcurver: an R package for obtaining interpretable metrics from microbial growth curves. BMC Bioinformatics 17: 172.

Stelkens, R. and Bendixsen, D.P. (2022) The evolutionary and ecological potential of yeast hybrids. Curr Opin Genet Dev 76: 101958.

Stelkens, R.B., Brockhurst, M.A., Hurst, G.D.D., Miller, E.L., and Greig, D. (2014) The effect of hybrid transgression on environmental tolerance in experimental yeast crosses. J Evol Biol 27: 2507–2519.

Strathern, J.N., Jones, E.W., Broach J. R, and Dujon, B. (1981) Mitochondrial genetics and functions. In Molecular Biology of the Yeast Saccharomyces Life Cycle and Inheritance. Strathern, J.N., Jones, E.W., and Broach, J.R. (eds). Cold Spring Harbor, NY: Cold Spring Harbor Laboratory Press, pp. 505–635.

Sun, S., Wang, B., Li, C., Xu, G., Yang, J., Hufford, M.B., et al. (2023) Unraveling Prevalence and Effects of Deleterious Mutations in Maize Elite Lines across Decades of Modern Breeding. Mol Biol Evol 40: msad170.

Van de Peer, Y., Mizrachi, E., and Marchal, K. (2017) The evolutionary significance of polyploidy. Nat Rev Genet 18: 411–424.

Villarreal, P., Molinet, J., Braun-Galleani, S., and Cubillos, F.A. (2025) Non-Conventional Yeasts as a Source of Genetic Diversity and Biotechnological Potential. Annu Rev Microbiol 79: 595–614.

Voordeckers, K., Kominek, J., Das, A., Espinosa-Cantú, A., De Maeyer, D., Arslan, A., et al. (2015) Adaptation to high ethanol reveals complex evolutionary pathways. PLoS Genet 11: e1005635.

Wang, S.A. and Bai, F.Y. (2008) *Saccharomyces arboricolus* sp. nov., a yeast species from tree bark. Int J Syst Evol Microbiol 58: 510–514.

Weiss, C.V., Roop, J.I., Hackley, R.K., Chuong, J.N., Grigoriev, I.V., Arkin, A.P., et al. (2018) Genetic dissection of interspecific differences in yeast thermotolerance. Nat Genet 50: 1501–1504.

Wickham, H. (2009) ggplot2: elegant graphics for data analysis, NY: Springer.

Wolfe, K.H. (2015) Origin of the Yeast Whole-Genome Duplication. PLOS Biol 13: e1002221.

Wu, X., Chen, X., Wang, R., Wang, H., Hu, X., Wang, Y., et al. (2025) Transcriptome dynamics and allele-specific regulation underlie wheat heterosis at the anthesis and grain-filling stages. BMC Genomics 26: 798.

Zavaleta Vasni, Pérez-Través Laura, Saona Luis A., Villarroel Carlos A., Querol Amparo, and Cubillos Francisco A. (2024) Understanding brewing trait inheritance in de novo Lager yeast hybrids. mSystems 0: e00762-24.

