## Supplementary Figures for "Mitochondrial inheritance as an important parameter for the rational design of synthetic yeast polyploids"

#### Mitochondrial inheritance as a critical parameter for the rational design of synthetic *Saccharomyces* allotetraploids

Sara Orellana-Muñoz, Chloé Haberkorn, Bálint Csoboz, Rike Stelkens, David Peris*

### Supplementary Figures

**Supplementary Figure 1. Phenotype of the *rho⁰* yeast strains.**

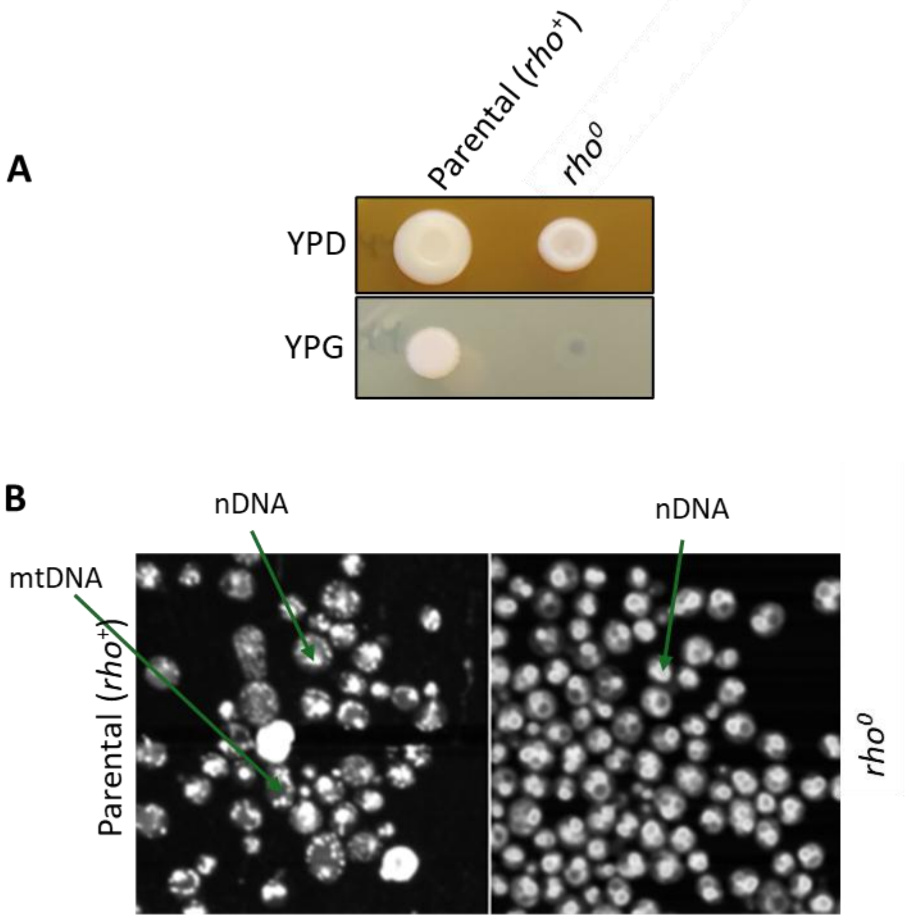

**A**. The parental strain *rho*$^{+}$ and its *rho*$^{0}$ version were grown on YPD and YPG at 20 °C. **B**. DAPI staining of both nuclear and mitochondrial DNA in the parental *rho*$^{+}$ and *rho*$^{0}$ strains. The DNA location was viewed by fluorescence microscopy.

**Supplementary Figure 2. Confirmation of newly synthetized allotetraploid strains.**

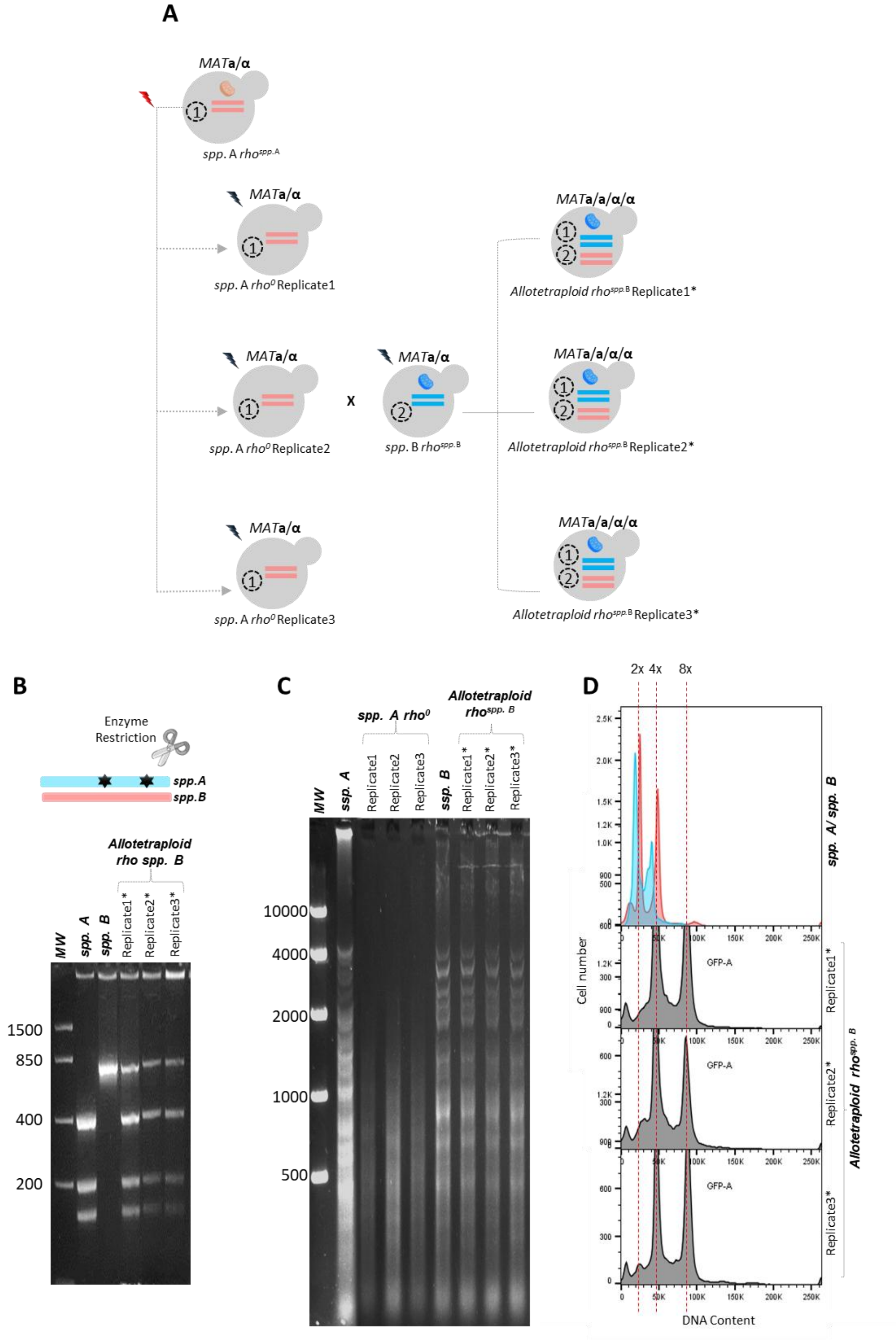

**A.** Synthetic allotetraploid generation scheme. 3 replicates per *rho*$^0$ strains and 3 replicates per allotetraploid cells (marked with an asterisk, "*"). **B.** RFLP (Restriction Fragment Length Polymorphism). Gene: *BRE5;* enzyme restriction: *Hae*III. **C.** mtDNA restriction pattern (*Hinf*I). *spp. A*: *S. paradoxus* (blue), *spp. B*: *S. arboricola* (pink). Allotetraploid: *S. paradoxus* x *S. arboricola.* **D.** The flow cytometry of green fluorescent protein (GFP-A) expression in both diploid parental strains (blue and pink distributions) and allotetraploid cells (gray distributions). The horizontal axis is the intensity of SYTOX Green fluorescence (DNA content). MW: Molecular weight ladder.

**Supplementary Figure 3. Positive correlation between DNA content (SYTOX Green) and cell size (FCS-A).**

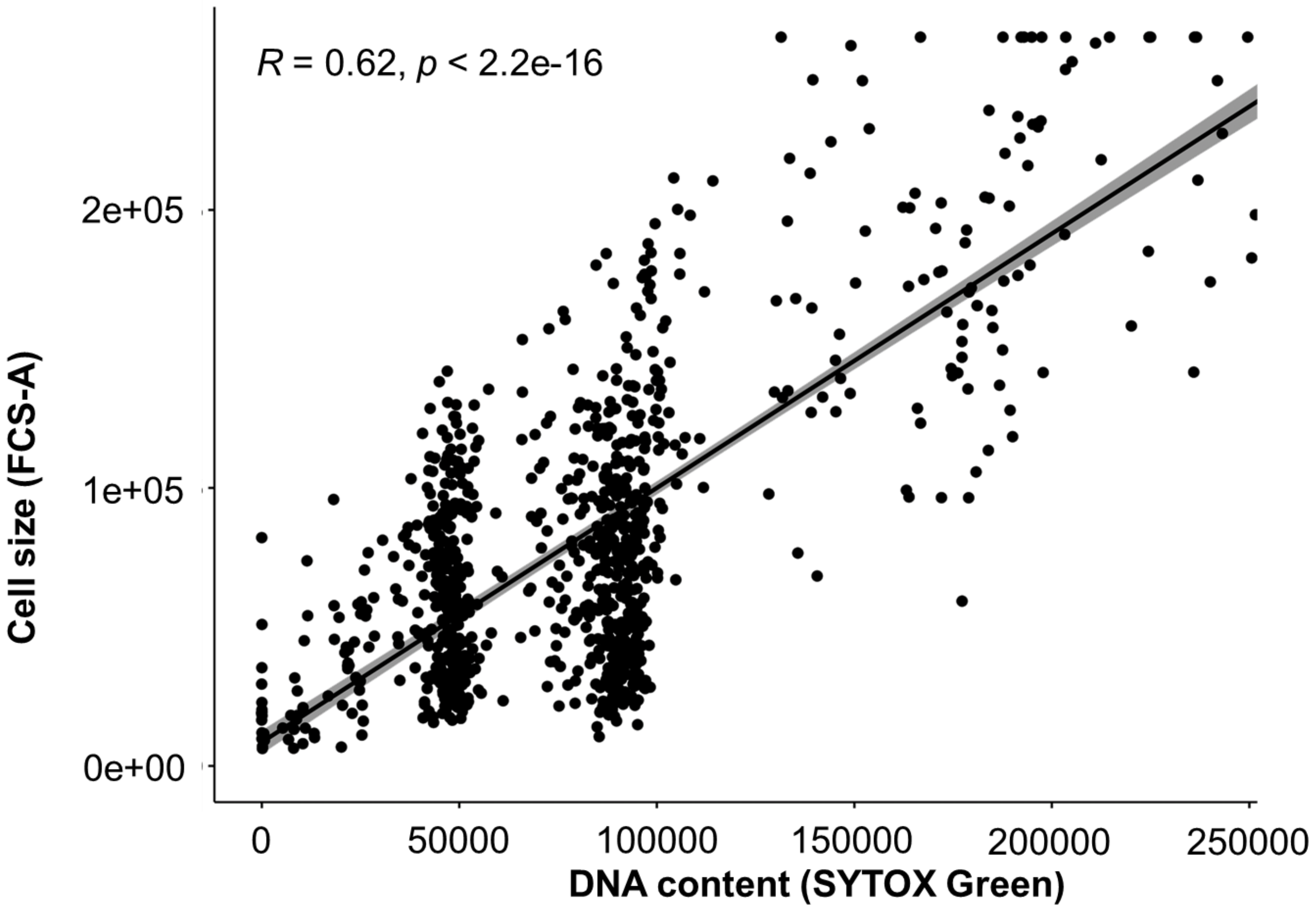

Spearman rank-order correlation coefficient (R) of SYTOX Green and FCS-A, together with the *p*-value are displayed. Allotetraploid example: *S. cerevisiae* x *S. paradoxus*.

**Supplementary Figure 4. Differential impact of mitochondrial DNA loss on growth across *Saccharomyces* species.**

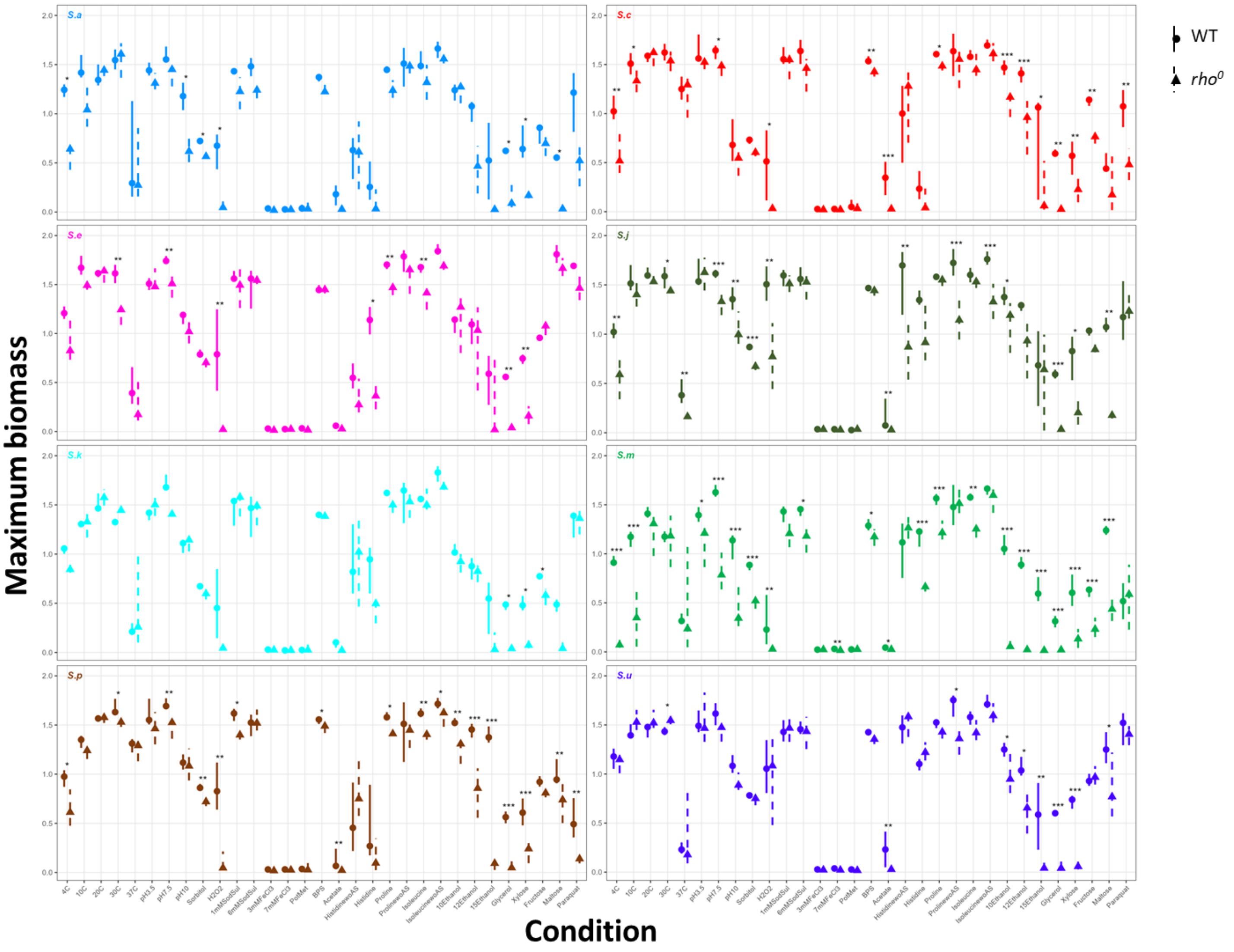

Median maximum cell growth and the upper and lower whiskers of the maximum and minimum values within 1.5× the interquartile range (IQR), respectively, were represented for the eight *Saccharomyces* species in wild-type (*rho*$^{+}$) and mitochondrial DNA-deficient (*rho*$^{0}$) backgrounds across 31 tested conditions.

**Supplementary Figure 5. Environmental conditions revealing mitotype-dependent phenotypic variation in synthetic allotetraploids.**

A

10% Ethanol

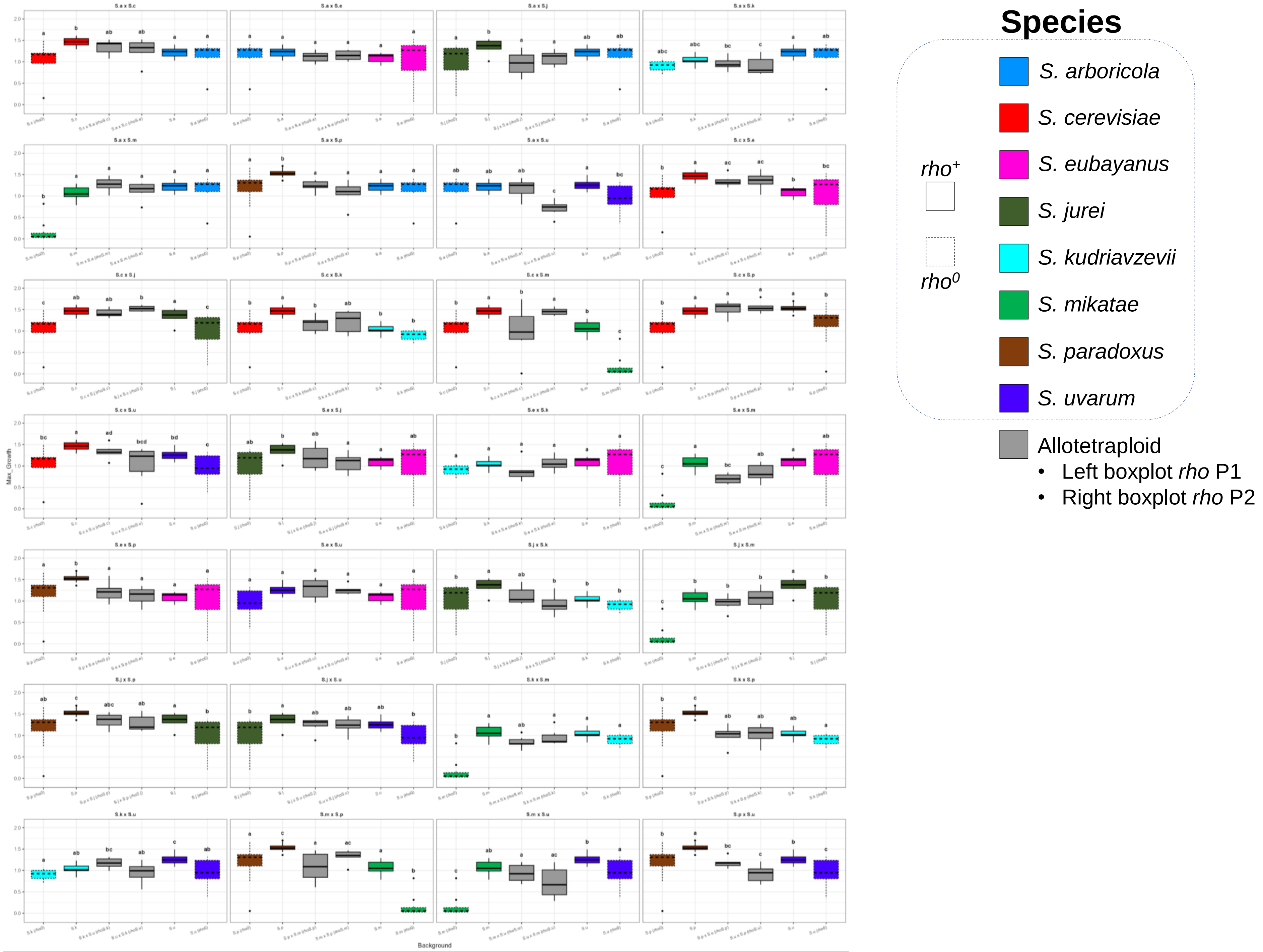
Species
S. arboricola
S. cerevisiae
S. eubayanus
S. jurei
S. kudriavzevii
S. mikatae
S. paradoxus
S. uvarum
rho+
rho0
Allotetraploid
• Left boxplot rho P1
• Right boxplot rho P2
Max_Growth
Background

**B**

12% Ethanol

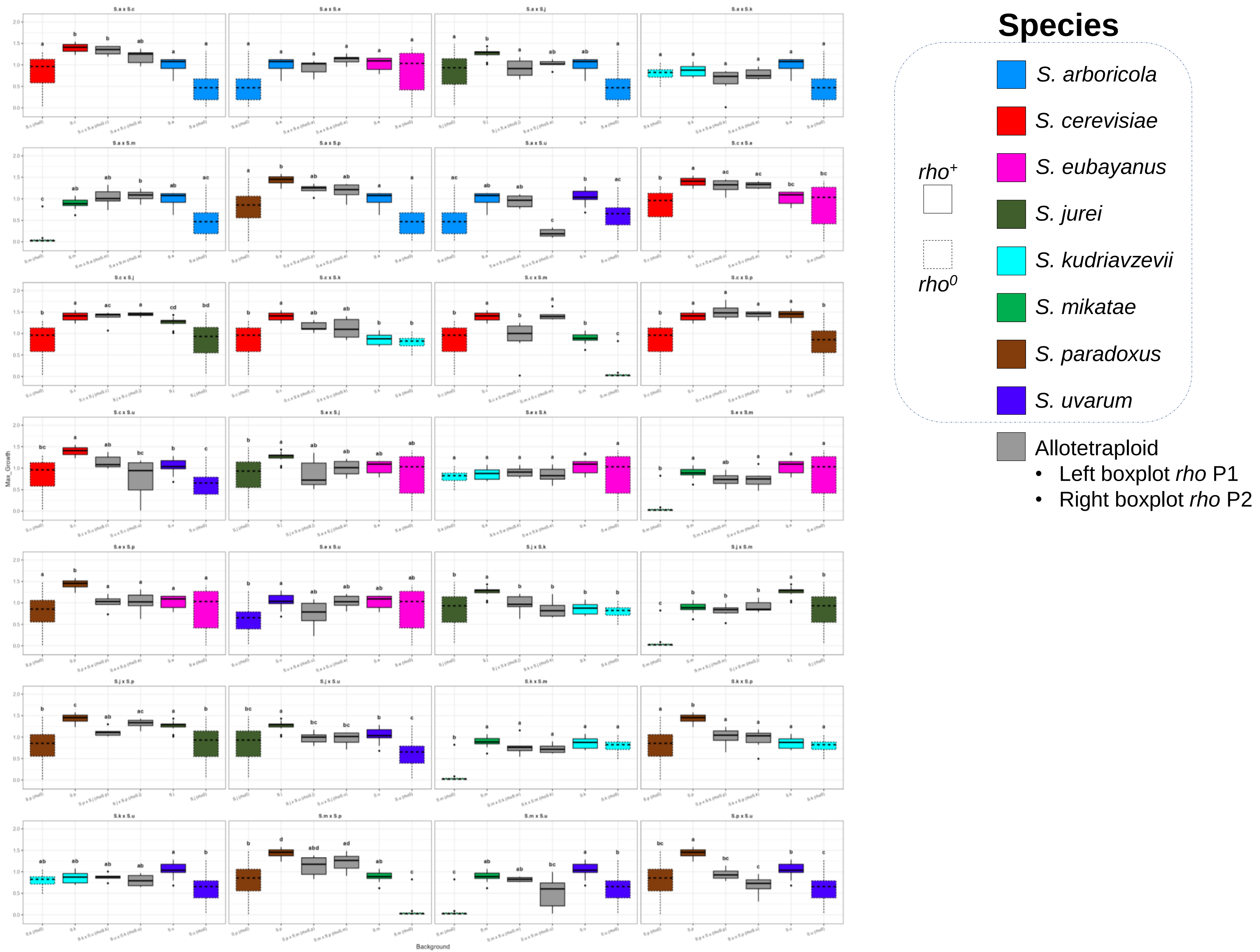

Species
rho+
rho0
S. arboricola
S. cerevisiae
S. eubayanus
S. jurei
S. kudriavzevii
S. mikatae
S. paradoxus
S. uvarum
Allotetraploid
• Left boxplot rho P1
• Right boxplot rho P2
Max_Growth
Background

C

15% Ethanol

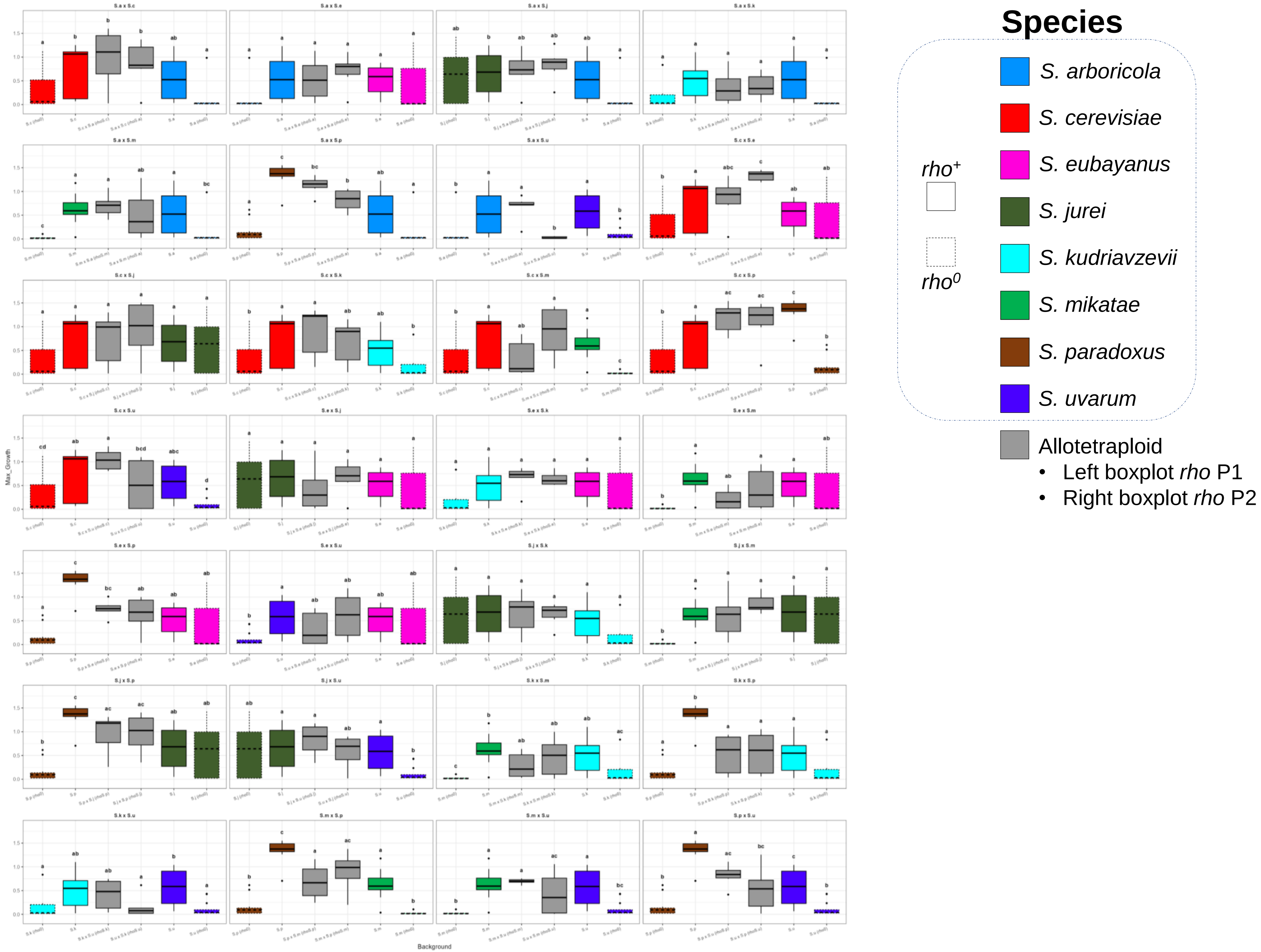

Species
$rho^{+}$
$rho^{0}$
S. arboricola
S. cerevisiae
S. eubayanus
S. jurei
S. kudriavzevii
S. mikatae
S. paradoxus
S. uvarum
Allotetraploid
• Left boxplot rho P1
• Right boxplot rho P2
Max_Growth
Background

**D**

2mM $H_2O_2$

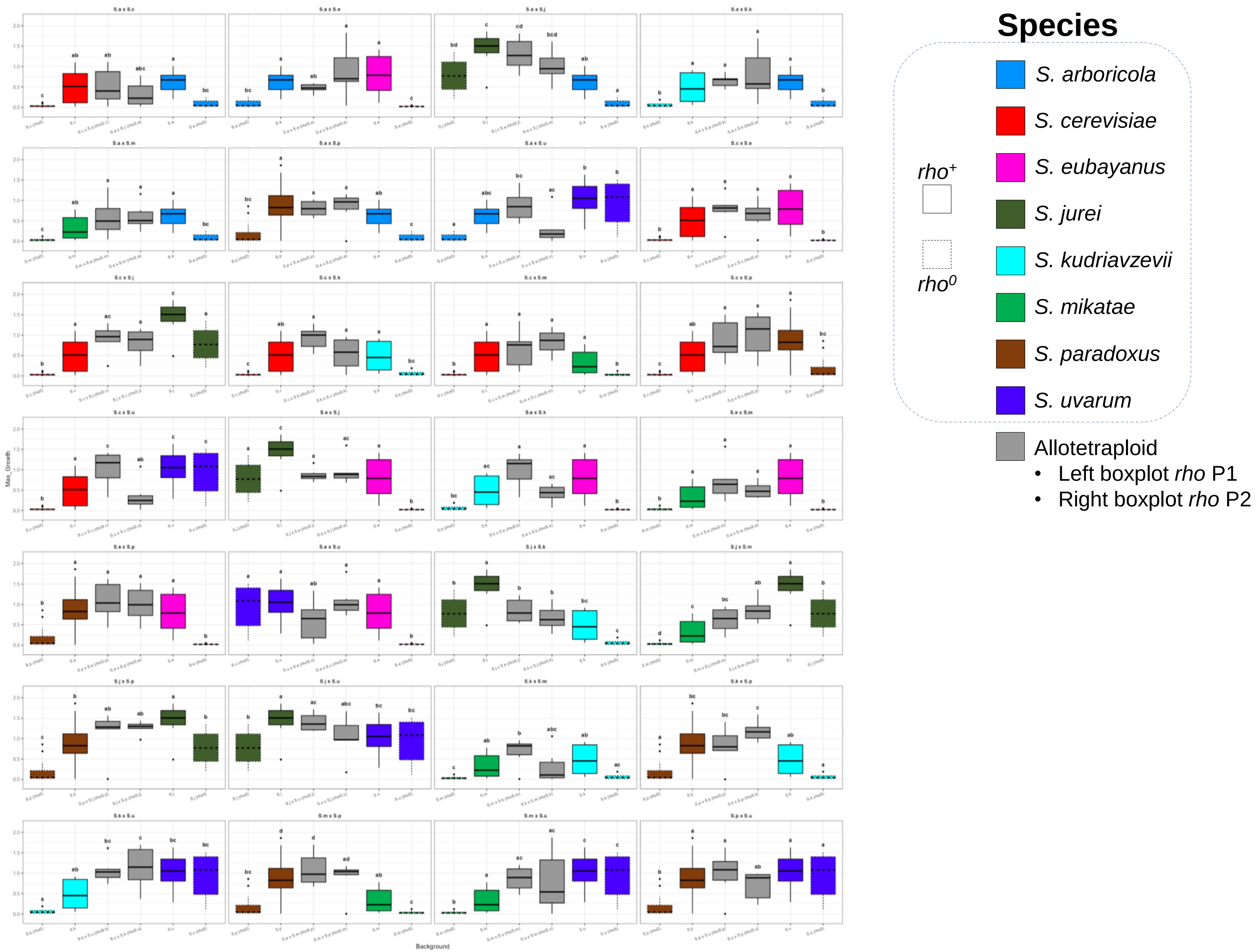
Max_Growth
Background
Species
*S. arboricola*
*S. cerevisiae*
*S. eubayanus*
*S. jurei*
*S. kudriavzevii*
*S. mikatae*
*S. paradoxus*
*S. uvarum*
*rho*+
*rho*0
Allotetraploid
• Left boxplot *rho* P1
• Right boxplot *rho* P2

**E**

200 µg/mL
Paraquat

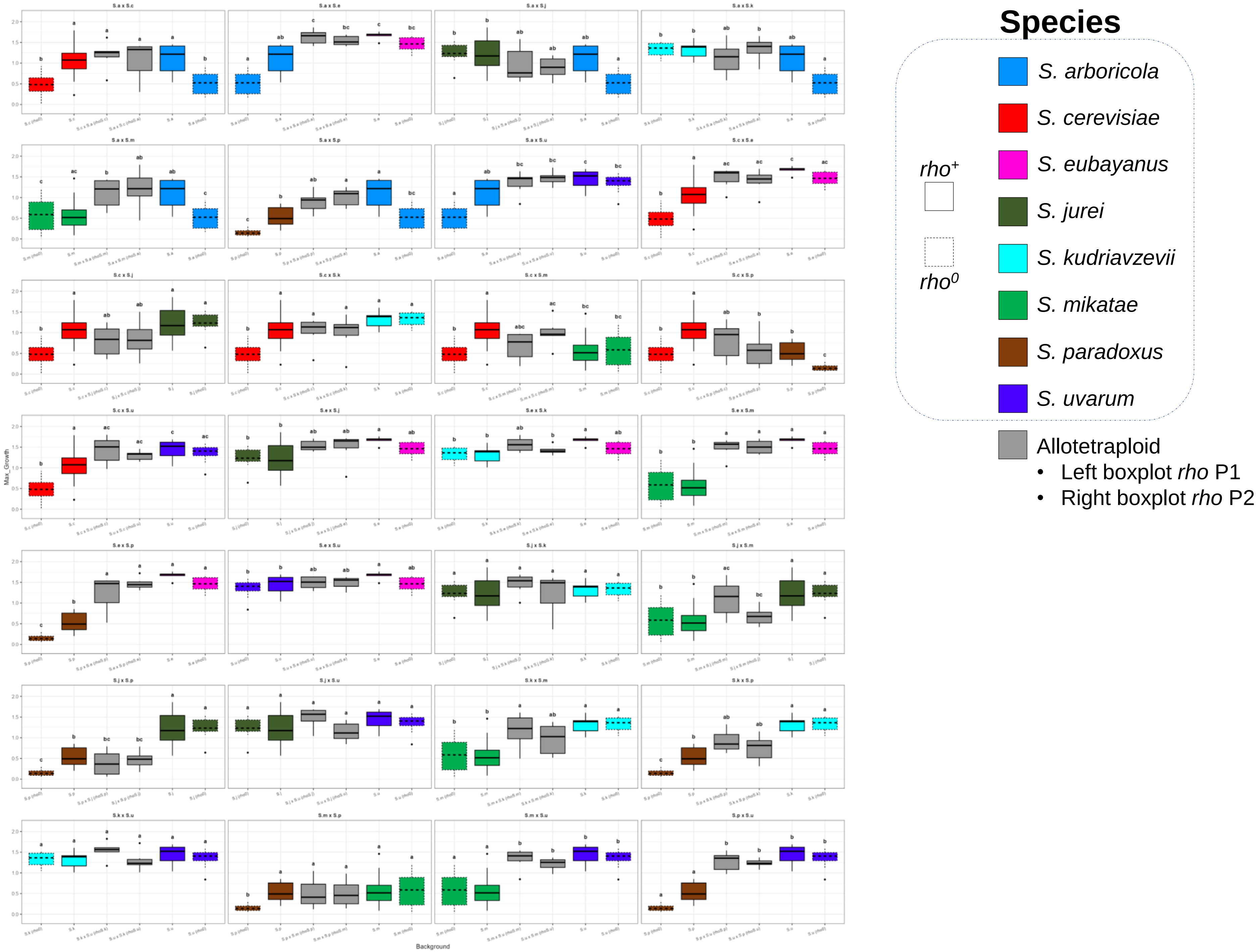

**F**

5g/L Histidine

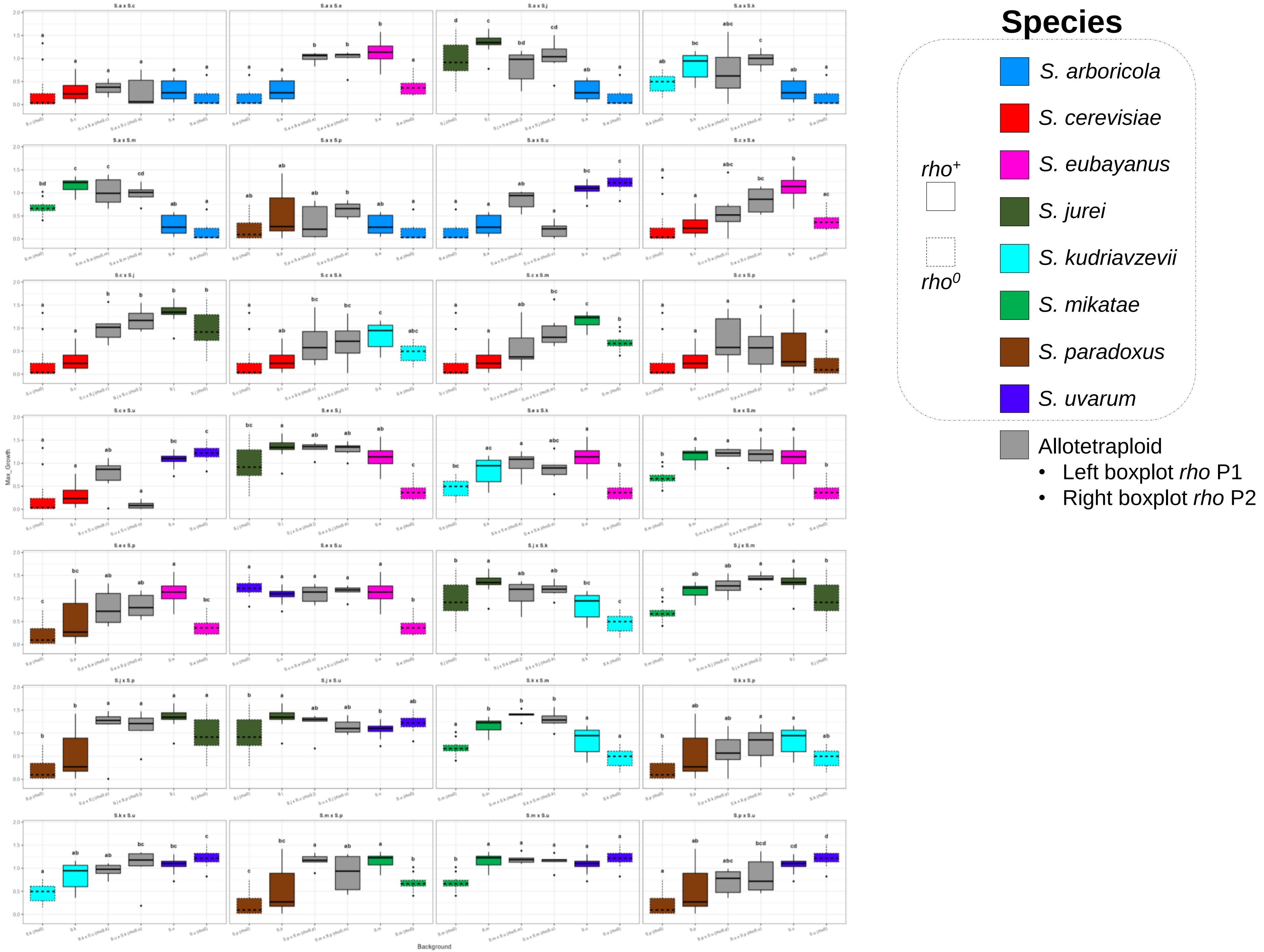

G

5.86 g/L Histidine
w/o AS

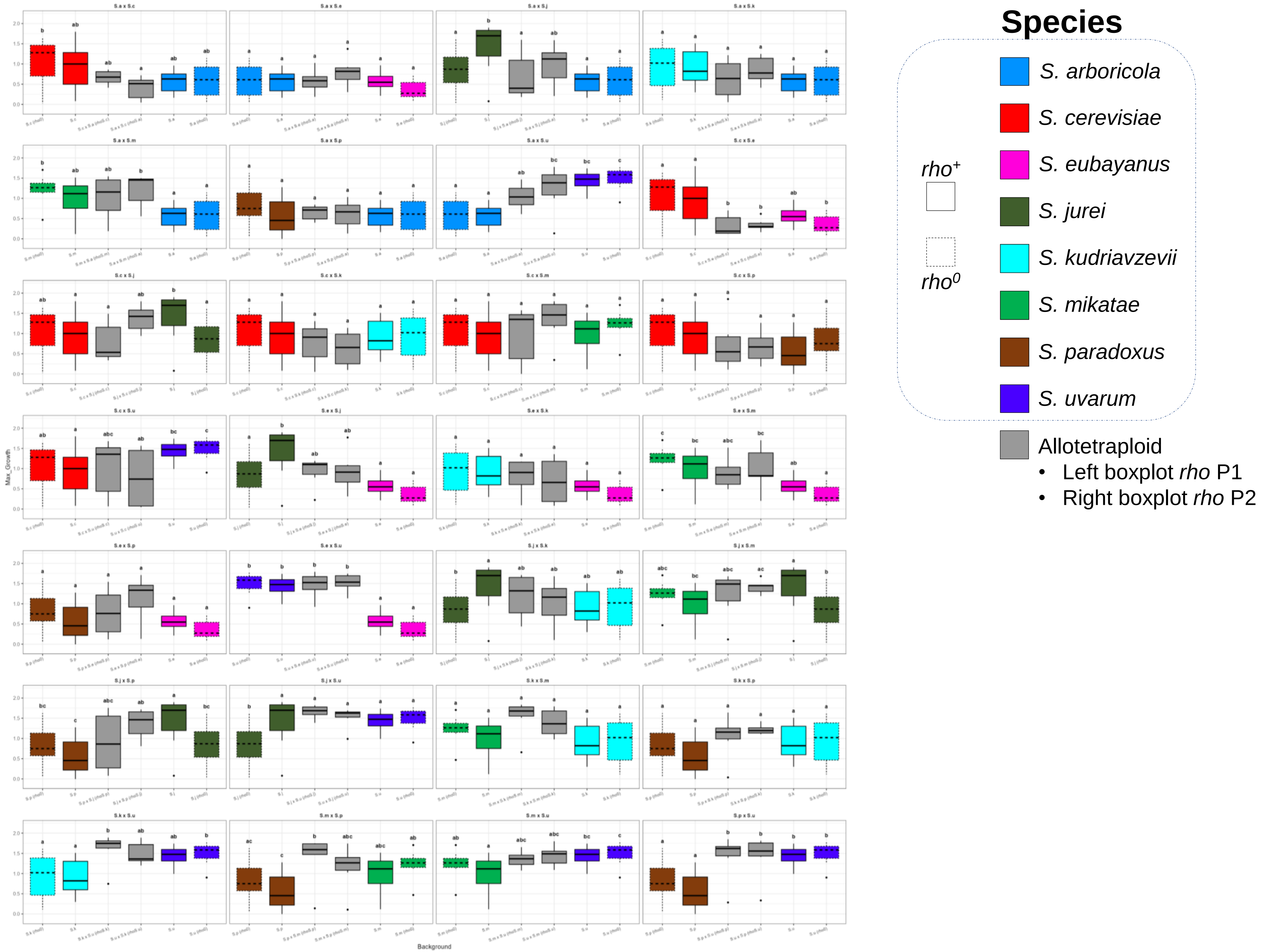

**H**

5g/L Proline
w/o AS

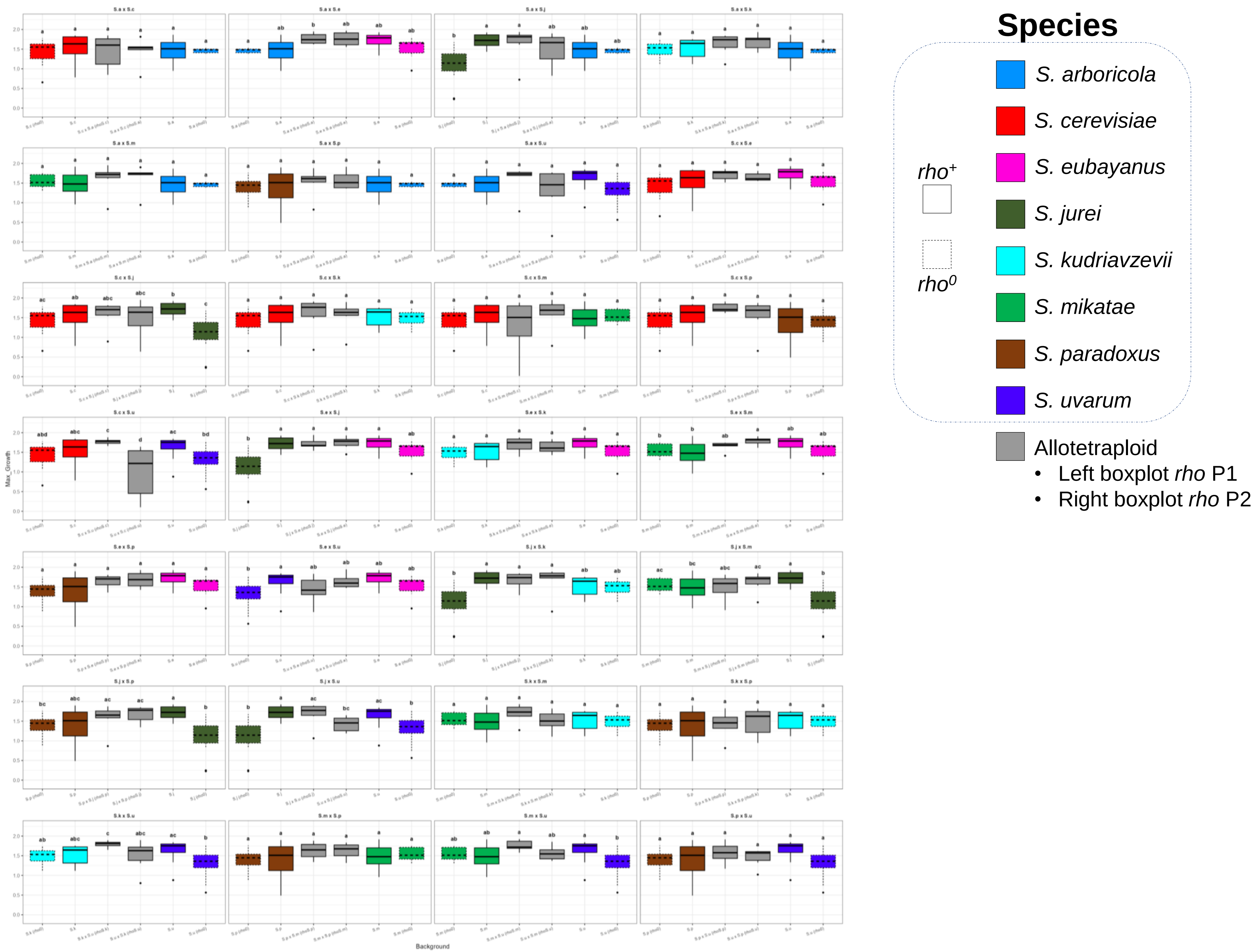

I

37°C

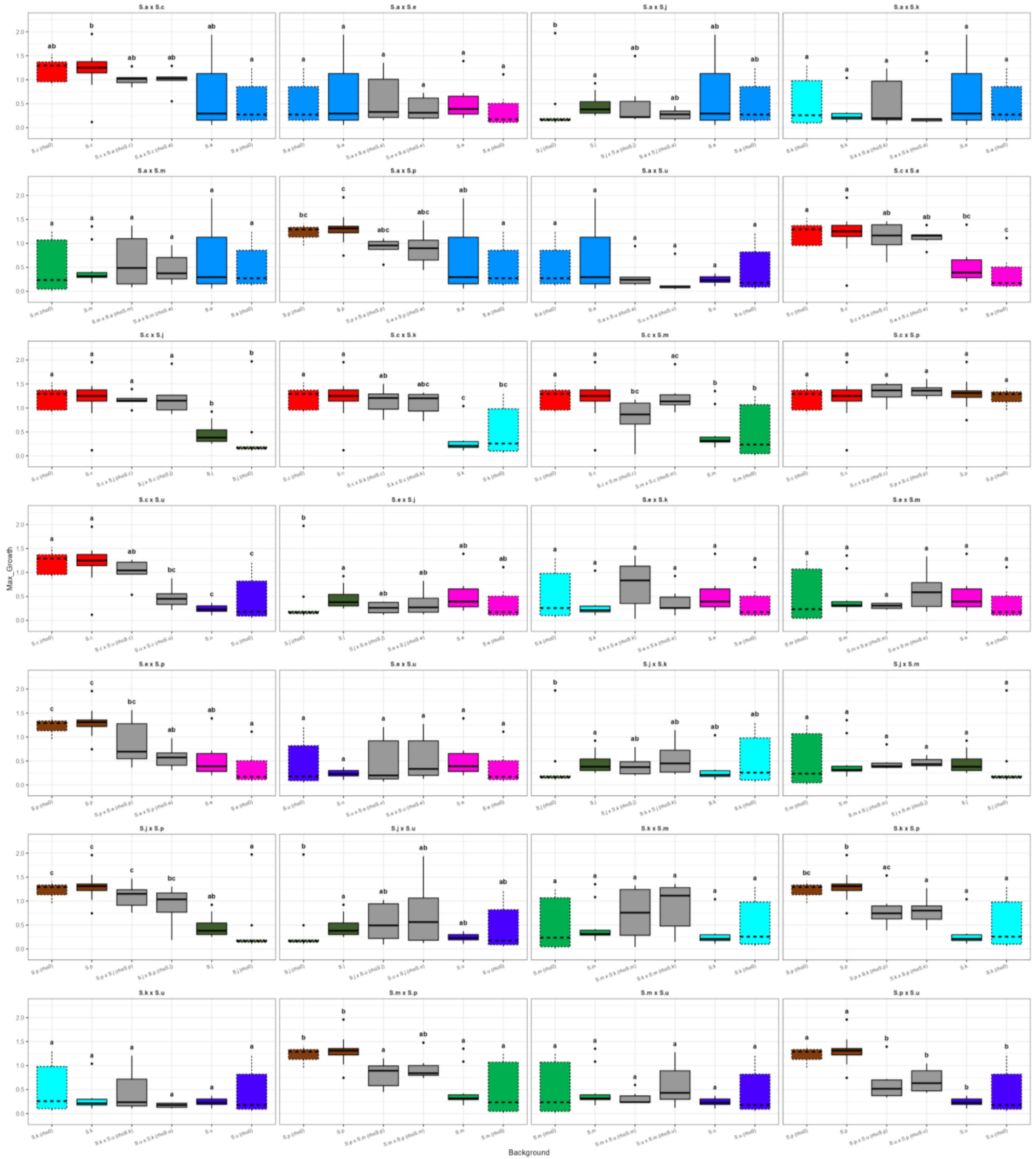

Species
S. arboricola
S. cerevisiae
S. eubayanus
S. jurei
S. kudriavzevii
S. mikatae
S. paradoxus
S. uvarum
rho+
rho0
Allotetraploid
• Left boxplot rho P1
• Right boxplot rho P2
Background

J

2% Maltose

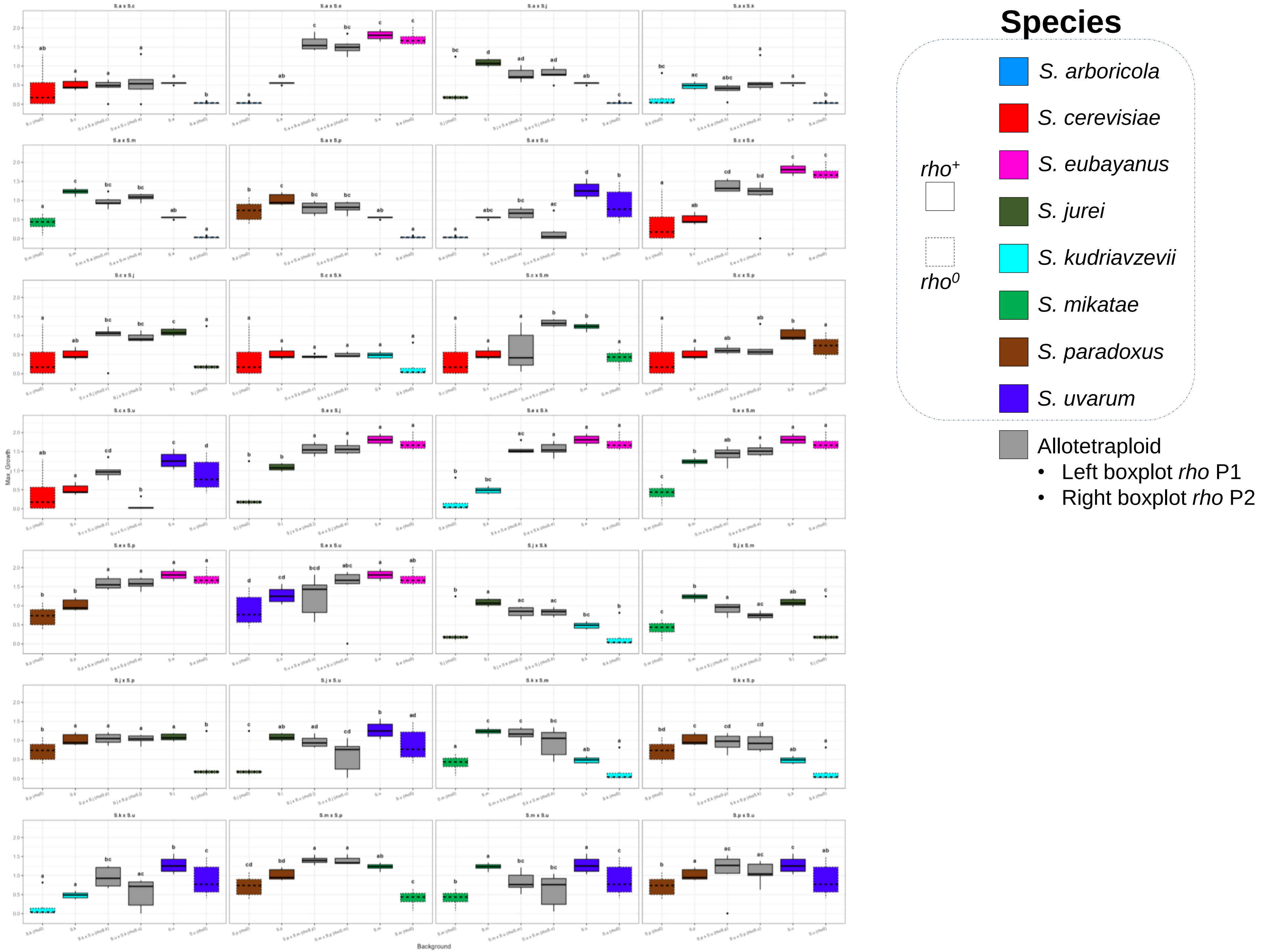
Species
S. arboricola
S. cerevisiae
S. eubayanus
S. jurei
S. kudriavzevii
S. mikatae
S. paradoxus
S. uvarum
rho+
rho0
Allotetraploid
• Left boxplot rho P1
• Right boxplot rho P2
Max_Growth
Background

Boxplots showing the distribution of maximum biomass for each genomic background. Each allotetraploid is presented alongside its corresponding parental strains, with and without (dashed boxplot) mitochondrial DNA, as indicated in the legend. Boxplots are colored according to the legend. The upper and lower whiskers represent the maximum and minimum values within 1.5× the interquartile range (IQR), respectively. Different letters indicate significant differences among groups based on a Dunn test with Bonferroni–Holm correction. Panels **A–J** correspond to the conditions identified as significant in the LMM analysis (Supplementary Figure 7).

**Supplementary Figure 6. Best parent heterosis is limited among environmental conditions.**

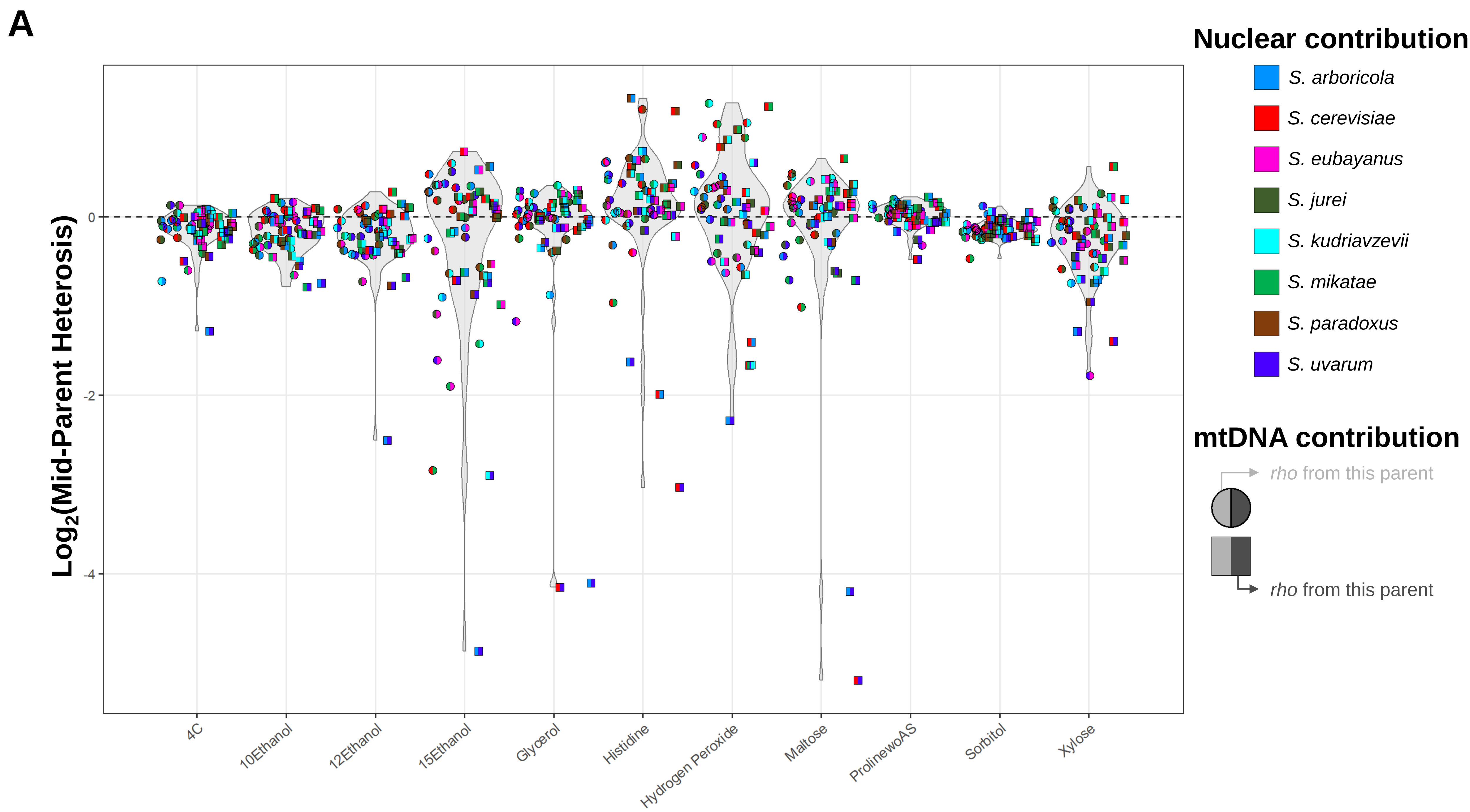
A
Nuclear contribution
S. arboricola
S. cerevisiae
S. eubayanus
S. jurei
S. kudriavzevii
S. mikatae
S. paradoxus
S. uvarum
mtDNA contribution
rho from this parent
rho from this parent
Log2(Mid-Parent Heterosis)
0
-2
-4
4C
10Ethanol
12Ethanol
15Ethanol
Glycerol
Histidine
Hydrogen Peroxide
Maltose
ProlinewoAS
Sorbitol
Xylose

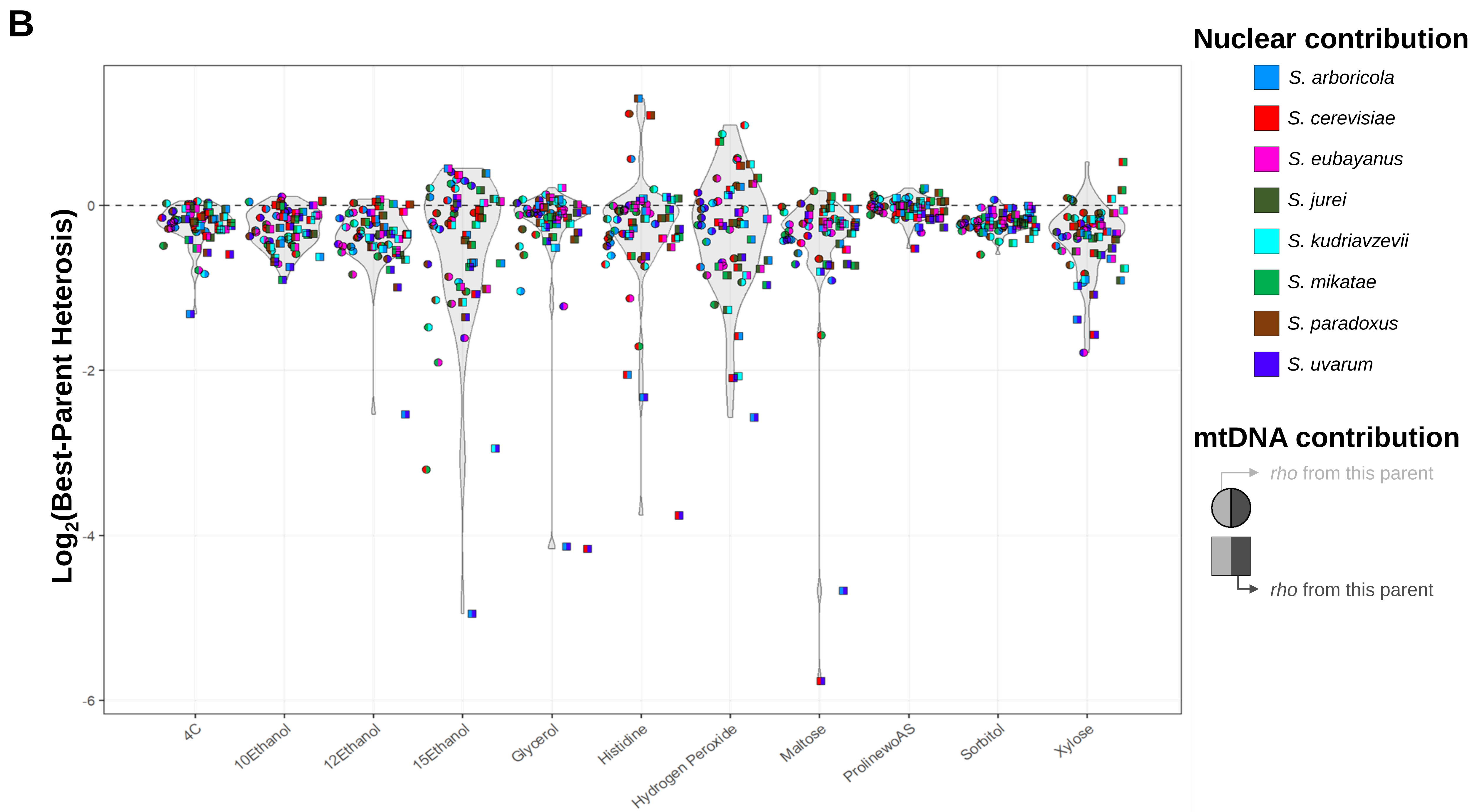
B
Log2(Best-Parent Heterosis)
0
-2
-4
-6
4C
10Ethanol
12Ethanol
15Ethanol
Glycerol
Histidine
Hydrogen Peroxide
Maltose
ProlinewoAS
Sorbitol
Xylose
Nuclear contribution
S. arboricola
S. cerevisiae
S. eubayanus
S. jurei
S. kudriavzevii
S. mikatae
S. paradoxus
S. uvarum
mtDNA contribution
rho from this parent
rho from this parent

Violin plots showing the distribution of **(A)** $\log_2$ Mid Parent Heterosis (MPH) calculate as [MPH = [(median Max_Growth of the hybrid - mean of the median Max_Growth P1 and median Max_Growth P2) / mean of the median Max_Growth P1 and median Max_Growth P2] or, **(B)** $\log_2$ Best Parent Heterosis (BPH), calculated as [BPH = [median Max_Growth of the hybrid / median Max_Growth of the best parent)] for each allotetraploid. Species contributions to each allotetraploid are represented as pie charts colored according to the legend. Mitotype inheritance is indicated by symbols: circles denote that the mitotype corresponds to the parent represented on the left side of the pie chart, whereas squares indicate that the mitotype derives from the parent represented on the right. Values above 0 denotes hybrid vigor (heterosis).

**Supplementary Figure 7. Mitochondrial DNA origin has an impact on allotetraploid growth across environments.**

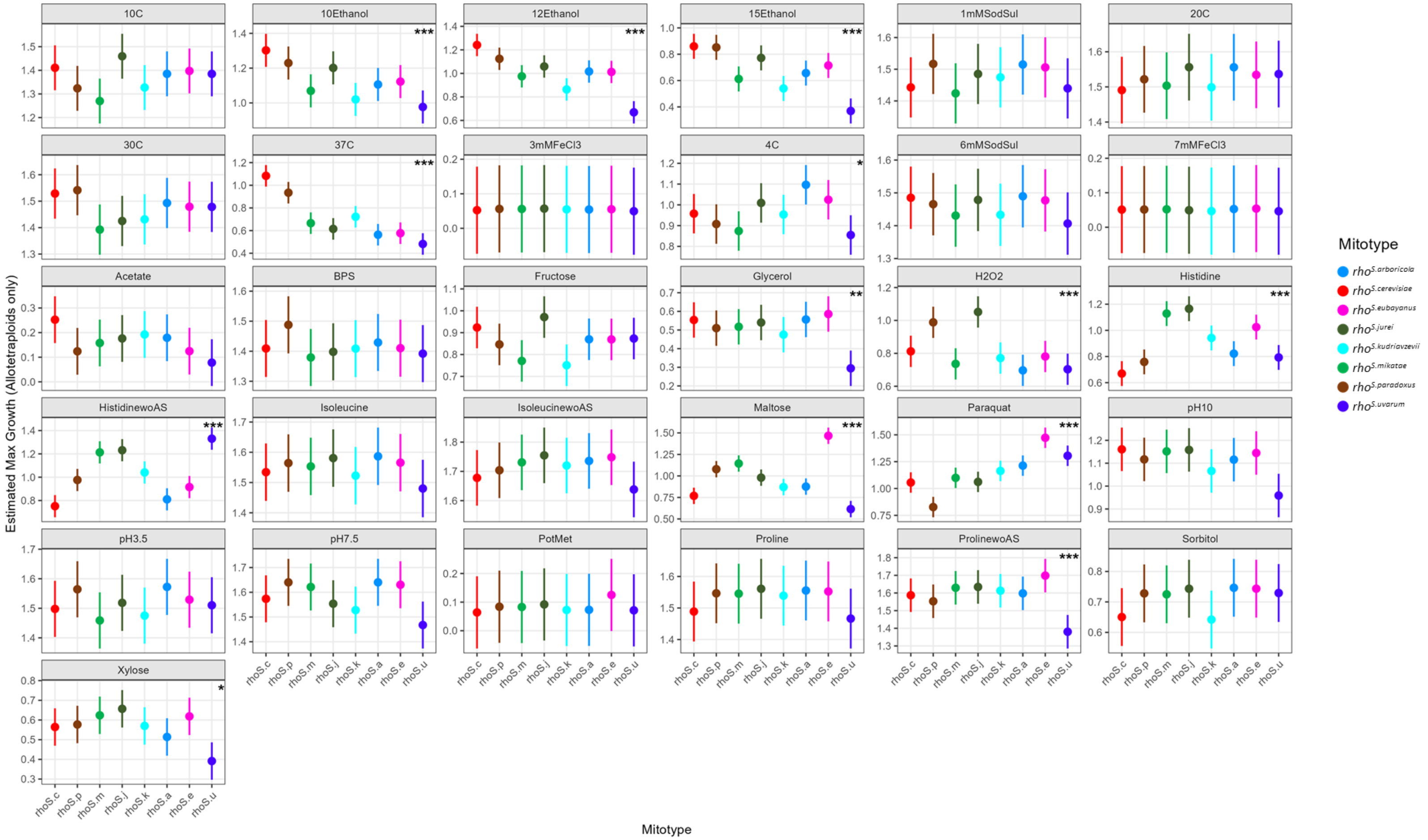

Estimated marginal mean (EMM) for maximum growth of allotetraploids bearing each mitotype, with 95% confidence intervals, for each condition. A higher EMM indicates that the model predicts greater growth for hybrids carrying a given mitotype under that specific condition. Stars denote significance of an omnibus test of the mitotype effect within each condition (*p*-value < 0.05 (*), *p*-value < 0.01 (**), *p*-value < 0.001 (***)); 10 of 13 significant conditions show p-value < 0.001. For the most significant conditions, maximum growth per condition for parents and allotetraploids, along with pairwise contrasts between mitotypes, is shown in Supplementary Figure 5. Note that y-axis scales differ across condition plots for improved visibility.
